# MiR-146a-dependent regulation of CD24/AKT/β-catenin axis drives cancer stem cell phenotype in oral squamous cell carcinoma

**DOI:** 10.1101/429068

**Authors:** Sangeeta Ghuwalewala, Dishari Ghatak, Sumit Das, Pijush Das, Ramesh Butti, Mahadeo Gorain, Gopal C Kundu, Susanta Roychoudhury

**Affiliations:** Cancer Biology and Inflammatory Disorder Division, CSIR-Indian Institute of Chemical Biology,4, Raja S.C. Mullick Road, Jadavpur, Kolkata-700032, India.; Laboratory of Tumor Biology, Angiogenesis and Nanomedicine Research, National Centre for Cell Science (NCCS), Pune 411007, India.; Division of Research, Saroj Gupta Cancer Centre and Research Institute, Mahatma Gandhi Road, Thakurpukur, Kolkata 700063, India

**Keywords:** miR-146a, β-catenin, Wnt, signaling, p-AKT, stemness, CD44, CD24.

## Abstract

Cancer stem cells (CSCs) are known to potentiate tumor initiation and maintenance in Oral Squamous Cell Carcinoma (OSCC). Increasing evidences suggest that CD44^high^CD24^low^ population in OSCC are potential CSCs. MicroRNAs (miRNAs) have emerged as crucial players in tumor development. However, their role in maintenance of OSCC stem cells remains unclear. Here we report that CD44^high^CD24^low^ population within OSCC cells and primary HNSCC tumors have an elevated expression of miR-146a. Moreover, over-expression of miR-146a results in enhanced stemness phenotype by augmenting CD44^high^CD24^low^ population. We demonstrate that miR-146a induces stemness by stabilizing β-catenin with concomitant loss of E-cadherin and CD24. Interestingly, CD24 is identified as a novel functional target of miR-146a and ectopic expression of CD24 abrogates miR-146a driven potential CSC phenotype. Mechanistic analysis reveals that higher CD24 levels inhibit AKT phosphorylation leading to β-catenin degradation. Using stably expressing miR-146a/CD24 OSCC cell lines, we also validate that the miR-146a/ CD24/AKT loop significantly alters tumorigenic ability *in vivo*. Furthermore, we confirmed that β-catenin trans-activates miR-146a, thereby forming a positive feedback loop contributing to stem cell maintenance. Collectively, our study demonstrates that miR-146a regulate CSCs in OSCC through CD24-AKT-β-catenin axis.

**Highlights:**

- MiR-146a induces cancer stem cell characteristics in OSCC by targeting CD24
- CD24 abrogates miR-146a mediated stemness via β-catenin degradation in non-CSCs
- Akt/Wnt pathway is critical for sustenance of miR-146a driven potential CSCs
- The miR-146a/CD24/AKT loop significantly alters tumorigenic ability *in vivo*

## 1. Introduction

OSCC is the most prevalent form of head and neck cancers worldwide with more than 60% individuals diagnosed with advanced tumors[1]. The oral CSCs are held responsible for tumor aggressiveness leading to treatment failures, relapse and development of metastases [2, 3]. Over the past decade, epigenetic re-programming has emerged as a crucial mechanism of regulating cancer stem cell dynamics [4]. MiRNAs, a small ncRNAs of 20-22 nucleotides is a known epigenetic modulator, de-regulation of which may have critical roles in disease development [5],[6].They can act either as oncogenes or as tumor-suppressors depending upon the specific genes targeted. In-fact miRNA associated signatures are now considered for cancer specific diagnostic and prognostic purposes [7].

MiRNAs not only regulate primary cellular functions like proliferation, differentiation, migration and invasion, but also directly or indirectly influence CSC functions [8–11]. These are mostly attributed to altered signaling pathways including cell surface markers, pluripotency factors, chemo-resistance and epithelial-to-mesenchymal transition (EMT) markers[12].Role of miR-34a, miR-145a and miR-200bc family in regulating CD44, Oct4, Sox2, KLF4, Bmi1, Zeb1/2 and Notch1 has been well established[13–16]. MiR-146a is predominantly an onco-miR, which directly targets *IRAK1*, *traf6*, and *numb* genes in OSCC and imparts tumorigenicity [17].Emerging evidence on miR-146a suggests that it directs the self-renewal process in colorectal cancer stem cells by regulating Snail-β-catenin axis which also contributes to EMT[18]. High nuclear accumulation of β-catenin, along with lowering of E-cadherin is frequently associated with higher tumor grade and poor prognosis in various cancers[19]. Given the role of Wnt/β-catenin signaling in CSC maintenance and the miRNAs in regulating wnt pathway, understanding the step-wise regulation of its mediators is crucial [20–22].

Our recent study has characterized CD44^high^CD24^low^ cells as the potential CSC population in OSCC[2]. CD24, a small cell surface protein, was identified as a critical determinant of differentiation in hematopoietic cells and mammary epithelial cells[23]. Besides role in adhesion, cadherin switching or migration, CD24 is involved in diverse signaling networks, thus promoting oncogenesis or regression[24]. Although role of CD44 is well established[3], the involvement of CD24 in determining stemness is less explored, particularly in oral CSCs. In this study, we show that miR-146a imparts CD44^high^CD24^low^status to OSCC cells merely by targeting CD24. Changes in β-catenin pools seemed to be an important event in altering the afore-mentioned phenotypes downstream of miR-146a through AKT pathway. We propose that miR-146a/CD24/AKT/β-catenin axis influences the stemness characteristics of CD44^high^CD24^low^ population in oral cancer cells.

## 2. Materials and methods

### 2.1 Cell culture and transfection

Human OSCC cell lines SCC131, SCC084 and SCC036 were obtained from Dr. Sussane Gollin, University of Pittsburgh. These cells were maintained in 5% CO_2_ at 37°C in DMEM medium supplemented with 10% fetal bovine serum (FBS) and antibiotics (Life Technologies, Thermo Fisher Scientific Inc., MA, USA). The ATCC (American Type Culture Collection) oral cancer cell line, SCC25 was cultured in complete DMEM-F12 medium and 400ng/ml Hydrocortisone (Sigma Aldrich) under similar conditions. For transfection, Lipofectamine^TM^ 2000 (Invitrogen) was used in serum free medium. Transfected cells were harvested after 48 hrs or 72 hrs for over-expression or knockdown studies respectively.

### 2.2 Plasmid constructs, miRNA inhibitors and siRNAs

We obtained mir-146a and mir-146a SDM expressing pU61 construct from Dr. Nitai. P. Bhattacharjee (SINP, Kolkata). The TOP-flash/FOP-flash reporters, dnTCF4, Numb, and pcDNA3.1 empty vector along with miR-146a promoter LucA, LucB and mLucA Luciferase constructs were kind gifts from Muh-Hwa Yang (Taiwan). Human CTNNB1 expression plasmid deposited by Eric Fearon was purchased from Addgene (#16828). CD24 cDNA cloned into the pCDNA3.1 vector and the full-length 3′-UTR of CD24 cloned into pMIR (Ambion) were obtained from Heike Allgayer (University of Heidelberg, Germany). Anti-miR-146a (AM-34a) (ID: AM11030) were obtained from Ambion, CD24 siRNA (a pool of 3 target specific siRNAs) and scramble siRNA from Santa Cruz. CTNNB1 shRNA constructs (Addgene # 18803) were provided by Dr. Mrinal Kanti Ghosh, IICB, Kolkata. MiR-146a over-expression cassette was sub-cloned from pU61 into the pLKO.1 TRC vector (Addgene plasmid #10878). Packaging plasmids psPAX2 (Addgene plasmid # 12260) and pMD2.G (Didier Trono, Addgene plasmid# 12259) was used to generate the miR-146a over-expression lentiviral particles and target cells were infected following the manufacturer’s protocol. Stable transduced cells were selected by puromycin (Gibco) and over-expression efficiencies were verified by qRT-PCR and western blotting. CD24 was co-transfected and clones were selected by G418.

### 2.3 Quantitative Real Time PCR

TRIzol (Invitrogen, Thermo Fisher Scientific Inc., MA USA) method was used for isolation of total RNA as per manufacturer’s instructions. 250 ng of RNA were converted to cDNAs using stem-loop primers specific for reverse transcription of individual miRNAs[25]. MiRNA cDNAs were amplified with forward primers specific for individual miRNAs and a URP, with U6 snRNA as an endogenous reference control. For mRNA expression changes, protocol was similar to that described previously [2]. SYBR Green master mix (Roche, USA) was used to perform qRT-PCR in the 7500 Fast Real-Time PCR instrument (Applied Biosystems, USA). Fold change values (2^−ΔΔCT^) were calculated from average of three independent experiments. Primer sequences of genes, miRNA forward and loop primers are listed in Supplementary File 1.

### 2.4 Analysis of TCGA and NCI-60 datasets

RNA and miRNA-seq data were acquired for a total of 292 HNSCC tumor specimens from TCGA (The Cancer Genome Atlas) data portal (https://tcga-data.nci.nih.gov/tcga/). We first grouped top 25% of CD44 high and low expressing tumors and then further sub-grouped 25% of these tumors based on CD24 expression. These were designated as CD44^high^CD24^low^ and CD44^low^CD24^high^, wherein we checked the differential expression of miR-146a and calculated statistical significance using R Limma Package. Node status of these patients was also correlated with miR-146a expression using GraphPad Prism5 software. NCI-60 miRNA expression dataset (GEO accession number GSE26375) was analyzed to compare the miR-146a expression between the epithelial and mesenchymal groups as classified earlier [26] using Mann Whitney’s u test.

### 2.5 Flow cytometry

CD44-PE and CD24-FITC (BD Pharmingen) conjugated antibodies were used for double staining of miR-146a transfected cells. Cells were then washed and subjected to flow cytometry on the BD LSRFortessa and analysed using BD FACSDiva 6.2 software. Isotype controls were included for the non-specific staining.

### 2.6 Sphere forming assay

MiR-146a transfected cells were trypsinised and a single cell suspension was ensured. Low attachment 6-well plate were used for re-seeding the cells at a density of 5000 cells/ml in DMEM-F-12 serum free media containing 1% B27 supplement, 20 ng/ml of EGF and 20 ng/ ml of bFGF (Invitrogen). 500 µl of media was added every 2-3 days. Photographs of the spheres were taken under inverted microscope (Leica TCS SP8; Germany) with 20X magnification at 7-14 days. All experiments were done in biological triplicates.

### 2.7 Immuno-fluorescence

Sorted populations of SCC131 were grown on cover-slips overnight and then fixed with chilled aceto-methanol (1:1). 0.03% Saponin (Calbiochem, Germany) was used for permeabilization followed by blocking with 3% BSA. Rabbit monoclonal antibody against β-catenin and mouse monoclonal antibody against CD24, CD44 (Cell signaling technology) were added at a dilution of 1:200 and incubated overnight. It was then probed with anti-rabbit-FITC and anti-mouse Alexa-Flour 633nm conjugated secondary antibody (molecular probes) and counter stained with DAPI (Invitrogen) for nuclear staining. Images were taken under a confocal microscope (Andor Spinning Disc Confocal Microscope, Andor Technology, Belfast, Ireland) at 60X magnification.

### 2.8 Western blotting

Cell lysates were prepared after 48 hrs of transfection in NP-40 lysis buffer (Invitrogen) and protease inhibitor cocktail (1X). Equivalent amounts of denatured protein samples were subjected to SDS-PAGE (8%-10%), separated by size and transferred on to PVDF membrane (Millipore, Billerica, USA). Antibodies used for immuno-blotting were polyclonal β-catenin, E-cadherin, CD44 and CD24, Involucrin (Santa Cruz Biotechnology, CA, USA), polyclonal Oct4 and Sox2 (Abcam), polyclonal C-myc, Akt and phospho-Akt (Cell Signaling Technology, USA). Bands were obtained using ECL substrate (Thermo Scientific, USA) from HRP-conjugated secondary antibody (Sigma). Proteasome Inhibitor MG132 (Calbiochem) and Akt inhibitor LY294002 (Cell signaling Technology, USA) were both used at a concentration of 50 µM. Transfected cells were treated for 4 hours before harvesting. Band intensities of each protein were analyzed by ImageJ to obtain densitometric values for their quantification. These were normalized to β-actin for individual experimental sets and fold change calculated. All the histograms were expressed as means ± S.D. of three different experiments and p values computed in GraphPad Prism 5 (Student’s two tailed t test).

### 2.9 *In vivo* tumor xenograft experiments

Animal experiments were performed following guidelines of the institutional animal ethics committee of National Centre for Cell Science, Pune. All the animals were issued under the project IAEC/2012/B183. To investigate the effect of miR-146a overexpression on OSCC growth *in vivo*, 3×10^6^ empty vector- and microRNA overexpression construct-containing SCC084 cells were injected subcutaneously into the dorsal flanks of eight NOD/SCID male mice (18 weeks old) on left and right side respectively. When palpable tumors could be seen the mice were segregated into groups of four each. Mice in one of the groups were injected with 25 mg/kg of body weight of Quercetin (Sigma) on every alternate day for a period of 15 days. The experiment was terminated when the average miR-146a over-expressing SCC084 tumor volumes in the group which received no quercetin reached about 1200 mm^3^. At the termination of the experiment, the animals were sacrificed by CO_2_ asphyxiation and the tumors were collected for further analysis. Tumor diameters were measured each time the quercetin was injected and at the termination of the experiment using digital Vernier Caliper. Excised tumor tissues were weighed and then stored in RNAlater solution (ThermoFisher Scientific) in −20°C freezer. Tumor volumes were determined using the following formula: π/ 6[(d1×d2)^3/2^]; where d1 and d2 are two different diameters of a tumor. In another experiment, to investigate effect of simultaneous overexpression of miR-146a and CD24 on OSCC growth *in vivo*, 3×10^6^ empty vector- and miR-146a and CD24 overexpression constructs-containing SCC084 cells were injected subcutaneously into the dorsal flanks of four NOD/ SCID male mice (15 weeks old) on left and right side respectively. Tumors volumes were measured when palpable growth could be observed. The experiment was terminated when tumor volumes reached 1300 mm^3^. The animals were euthanized by CO_2_ asphyxiation and the tumors were collected. Tumor tissues were processed as described previously for the other experiment.

### 2.10 Reporter assays

Cells seeded in 24 well plates were co-transfected with miR-146a OE plasmid and either CD24 3′UTR or miR-146a promoter luciferase construct using Lipofectamine^TM^ 2000 (Invitrogen). The TOP-Flash and FOP-Flash reporters were also used under similar conditions. Promega dual luciferase assay system was performed according to the manufacturer’s protocol. After 48 hr of transfection, medium was washed off with 1x PBS and cells were lysed with Passive Lysis Buffer (Promega) and luminescence was measured in Promega Glomax 20/20 luminometer. The luminescence values were transfection normalized with the internal control pRL-TK (50 ng, Renilla Luciferase; Promega). Experiments were performed with three biological replicates.

### 2.11 Chromatin immunoprecipitation

Cells seeded in 10cm dishes were transfected. After 48hrs, 1X formaldehyde solution was added for DNA-protein crosslinking. Cells were lysed in SDS lysis buffer followed by sonication in Bioruptor (Diagenode) to obtain 200-1000 bp chromatin fragments. ChIP dilution buffer was used to dilute the sheared chromatin followed by preclearing with Protein G Agarose beads (Sigma) for 30min. After preclearing, 20% of the lysate was kept aside as the input and the remaining was divided equally for IP and IgG. Immunoprecipitation was carried out using 5µg of β-catenin (Santa Cruz) and normal IgG control (Sigma) and incubated overnight. The following day, Protein G Agarose beads were added to collect the Antibody/Antigen/Chromatin complex. The complex was washed briefly with cold low salt immune complex wash followed by high salt immune complex buffer, lithium chloride immune complex buffer and Tris-EDTA buffer. It was then reverse-crosslinked and the DNA purified using Phenol/Chloroform extraction method. PCR amplification of the immunoprecipitated DNA was carried out using primers listed in Supplementary File 1. Composition of the ChIP buffers are provided in Supplementary methods.

### 2.12 Statistical analyses

Three different experiments were subjected to an independent two-tailed Student’s *t* test to measure the significance value of results under varying biological parameters. R package was used to generate the correlation graphs and calculate p values. * indicates P ≤ 0.05 and ** indicates P ≤ 0.01.

## 3. Results

### 3.1 MiR-146a is over-expressed in CD44^high^CD24^low^ population of OSCC cell lines and primary tumors

To identify the cellular miRNAs regulating CSC phenotype of OSCC cells, we initially screened nine miRNAs that are aberrantly expressed in human cancers with their reported role in cancer stemness and EMT [27–29]. QRT-PCR data showed significant difference in the expression of miR-200b, miR-138, miR-34a, miR-21 and miR-146a betweenCD44^high^CD24^low^ and CD44^low^CD24^high^ population of SCC25 cells (Fig. 1a). Amongst them, we focused on miR-146a in view of its context dependent role in various cancers[30–32].MiR-146a is consistently over-expressed in oral CSCs, therefore it was intriguing to explore its possible connection with stemness and the underlying mechanisms. Up-regulation of miR-146a in the CD44^high^CD24^low^ population of SCC131 and SCC084 cell lines was also confirmed (Fig.1b). Further, miR-146a expression was also increased upon enrichment of isolated CSCs in the sphere forming culture conditions, suggesting its importance in determining oral CSC property (Fig. 1c). Increased miR-146a expression has been reported to predict poor survival of OSCC patients[17]. Our analyses in TCGA Head and Neck Squamous cell Carcinoma (HNSCC) patient’s cohort[33], showed increased miR-146a expression in patients with CD44^high^CD24^low^signature compared to those with CD44^low^CD24^high^profile (Fig. 1d). While there was not much difference in the histological stage of the tumors across the two categories, most of the CD44^low^CD24^high^ tumors were free of lymph node metastasis (Figure S1a, Supplementary file 1). Moreover, miR-146a expression of the node positive patients was relatively higher than that of the node negative ones, although not statistically significant (Figure S1b). Together, our data suggests high miR-146a expression as a critical determinant of CSC phenotype in oral tumors.

**Fig. 1.**
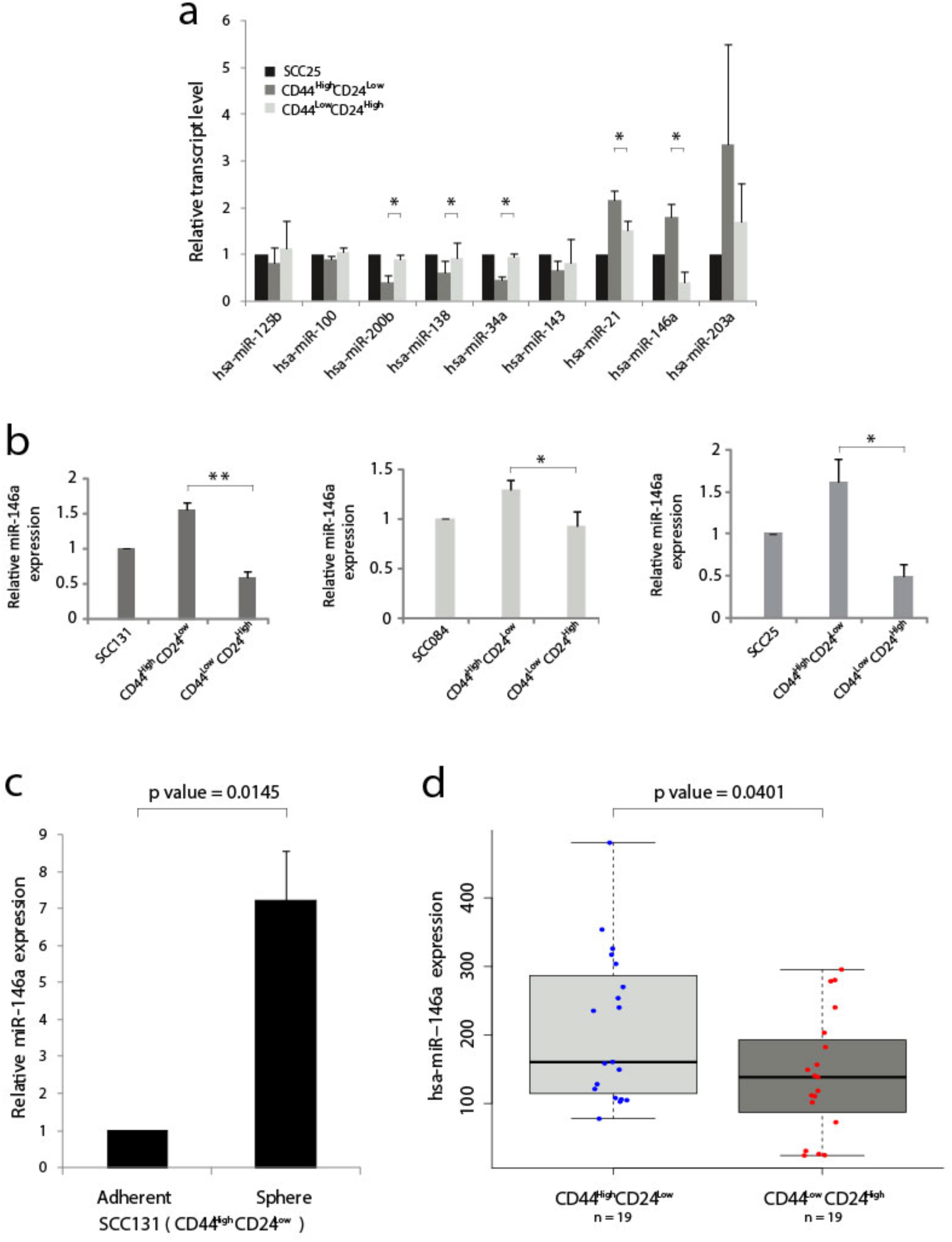
Over-expression of miR-146a in CD44^high^CD24^low^ cells of Oral Squamous Cell Carcinoma. (**a**) Total RNA extracted from the stem (CD44^high^CD24^low^) and the non-stem (CD44^low^CD24^high^) sub-populations of SCC25 cells were reverse-transcribed using stem-loop primers specific for subsequent Real time PCR analysis of various miRNAs using respective forward primers and Universal reverse primer (Supplementary File 1). (**b**) Expression of miR-146a was re-analyzed in UPCI: SCC131 and UPCI: SCC084 by qRT-PCR. (**c**) Quantification of miR-146a transcripts in the spheres enriched from CD44^high^CD24^low^ cells of SCC131 compared to that grown under differentiating adherent conditions. Data is representative of 3 independent experiments and the bar graph is shown with mean ± SD (right). U6snRNA was used to normalize relative expression values. Students t test used to calculate p-value (* P<0.05, **P<0.01, *** P<0.001). (**d**) Box-Scatter plot showing the differential expression of miR-146a in the CD44^high^CD24^low^and CD44^low^CD24^high^ subgroup of HNSCC tumors obtained from TCGA. (p-value was calculated in R package to show statistical significance).

### 3.2 Ectopic expression of miR-146a induces CSC characteristics

We next investigated whether ectopic miR-146a affect CSC markers and found a significant increase in the relative proportion of CD44^high^CD24^low^ population in SCC131 cells (Fig.2a). Similar results were also obtained with SCC036, SCC084 and SCC25 cells, respectively (Figure S2a). Characteristic sphere forming ability of miR-146a expressing SCC131 and SCC036 cells were also markedly enhanced (Fig. 2b). Concomitantly, ectopic expression of miR-146a led to the increased expression of intracellular stem cell markers such as Oct4, Sox2 and C-myc and loss of Involucrin (Fig. 2c). Also, we observed a pronounced decrease in CD24 protein levels upon ectopic miR-146a expression in SCC036, SCC131 and SCC084 cells (Fig. 2c). However, in SCC131 cells the levels of Oct4 and Involucrin did not show a dose dependent change upon miR-146a over-expression, probably due to high endogenous miR-146a in this cell line (Fig. 2c and Figure S2b). Loss of these markers upon knockdown of miR-146a was also evident in SCC131 cells (Figure S2c). Expression of miR-146a transcripts was validated by qRT-PCR (Figure S2d). Transfection of miR-146a containing mutated seed sequence, however, did not alter the levels of stem-related proteins in a statistically significant manner (Fig. 2d).These results demonstrate that miR-146a contributes to enrichment of CSCs in OSCC through increased expression of stem cell markers.

**Fig. 2.**
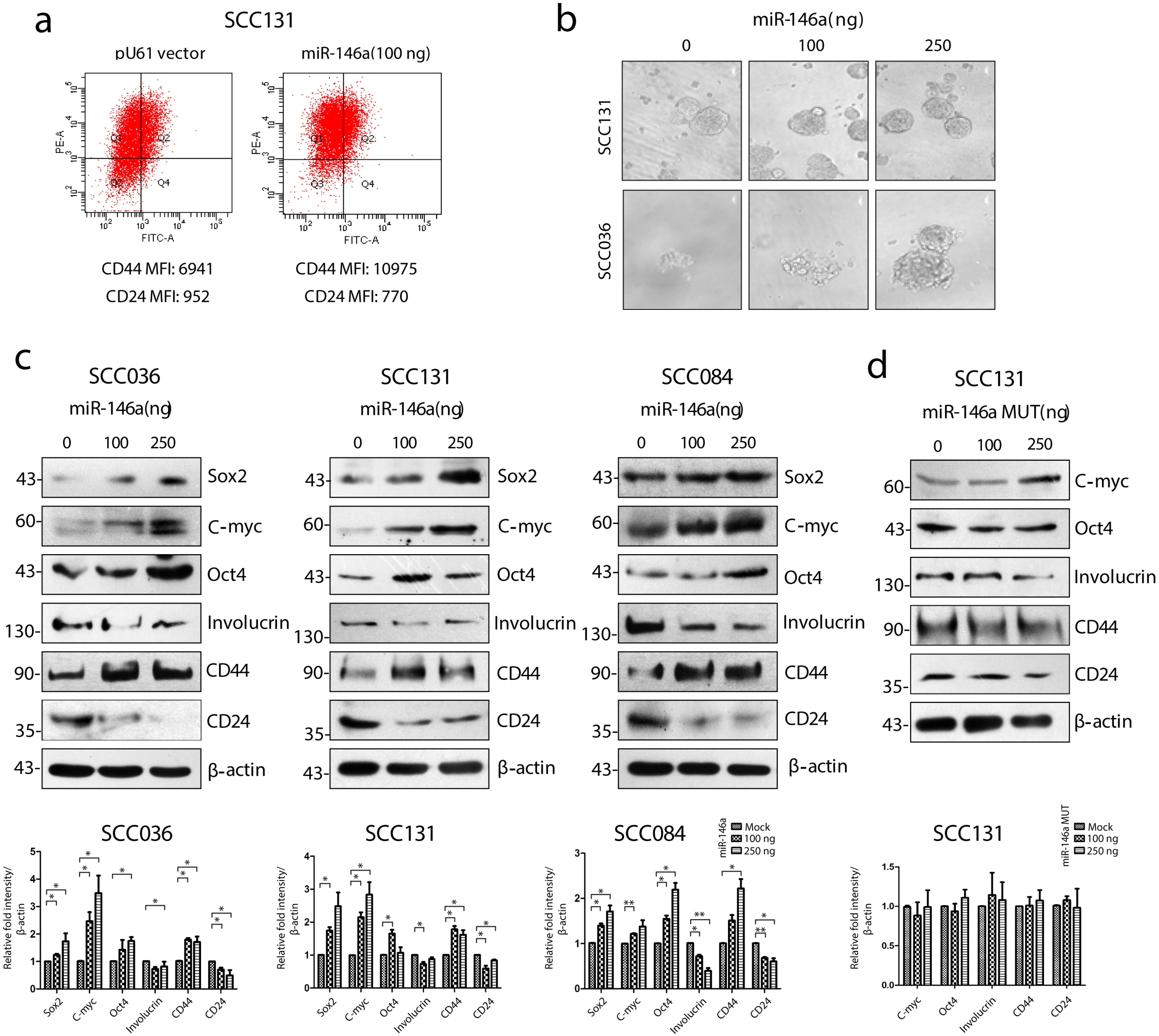
Cancer stem cell characteristics induced by miR-146a in OSCC cell lines. (**a**) MiR-146a transfected SCC131 cells were analyzed by flow cytometry and mean fluorescence values of CD44 (PE) and CD24 (FITC) are shown. (**b**) Equal number of vector and miR-146a transfected UPCI: SCC131 and UPCI: SCC036 cells were seeded in ultralow-attachment 6-well plate at clonal density. Sphere forming structures was captured at five random fields at 20X magnification using phase contrast microscope (Leica CTR4000) with scale bar equal to 50 µm. (**c**)Representative images of western blots showing dose dependent increment of Sox2, C-myc, Oct-4, Involucrin and CD44, CD24 in the UPCI: SCC036, UPCI: SCC131 and UPCI: SCC084 upon ectopic expression of miR-146a. β-actin bands was used to normalize the data. Band intensities of each protein has been quantified from three biological replicates and the average ± sd is plotted in GraphPad prism 5 showing statistical significance in the respective graphs (below)(* P<0.05, **P<0.01, *** P<0.001)(**d**) Similar western blots upon transfection of miR-146a with mutated seed sequences in UPCI:SCC131 and its graphical representation (below).

### 3.3 MiR-146a activates Wnt/β-catenin pathway and promotes EMT in oral CSCs

Wnt, Notch and Hedgehog pathways are often involved in maintaining stem cell self-renewal potential [34]. Accordingly, we observed increased expression of β-catenin and Cleaved Notch1 in CD44^high^CD24^low^ population of SCC25 and SCC131 cells (Figure S3a, b).While significantly depleted in CD24^high^cells, β-catenin levels were also higher in CD24^low^ population of SCC084 cells which correlated well with the expression of stemness markers in these cells(Fig.3a). To specifically envisage the role of CD24, we now compared the CD44^high^CD24^low^ population with the CD44^high^CD24^high^ cells only. Interestingly we detected nuclear localization of β-catenin in the CD44^high^CD24^low^ population of SCC131 cells, whereas it remained membrane bound in CD44^high^CD24^high^cells(Fig.3b). Transcriptional activity of β-catenin was indicated by the relative wnt reporter activity in the respective cell populations (Figure S3c). Interestingly, upon stable knockdown of β-catenin, not only the stem cell markers were reduced but also a modest increase in CD24 expression was observed (Figure S3d). These results suggest a possible cross talk between CD24 and β-catenin in conferring stemness and EMT to these cells.

**Fig. 3.**
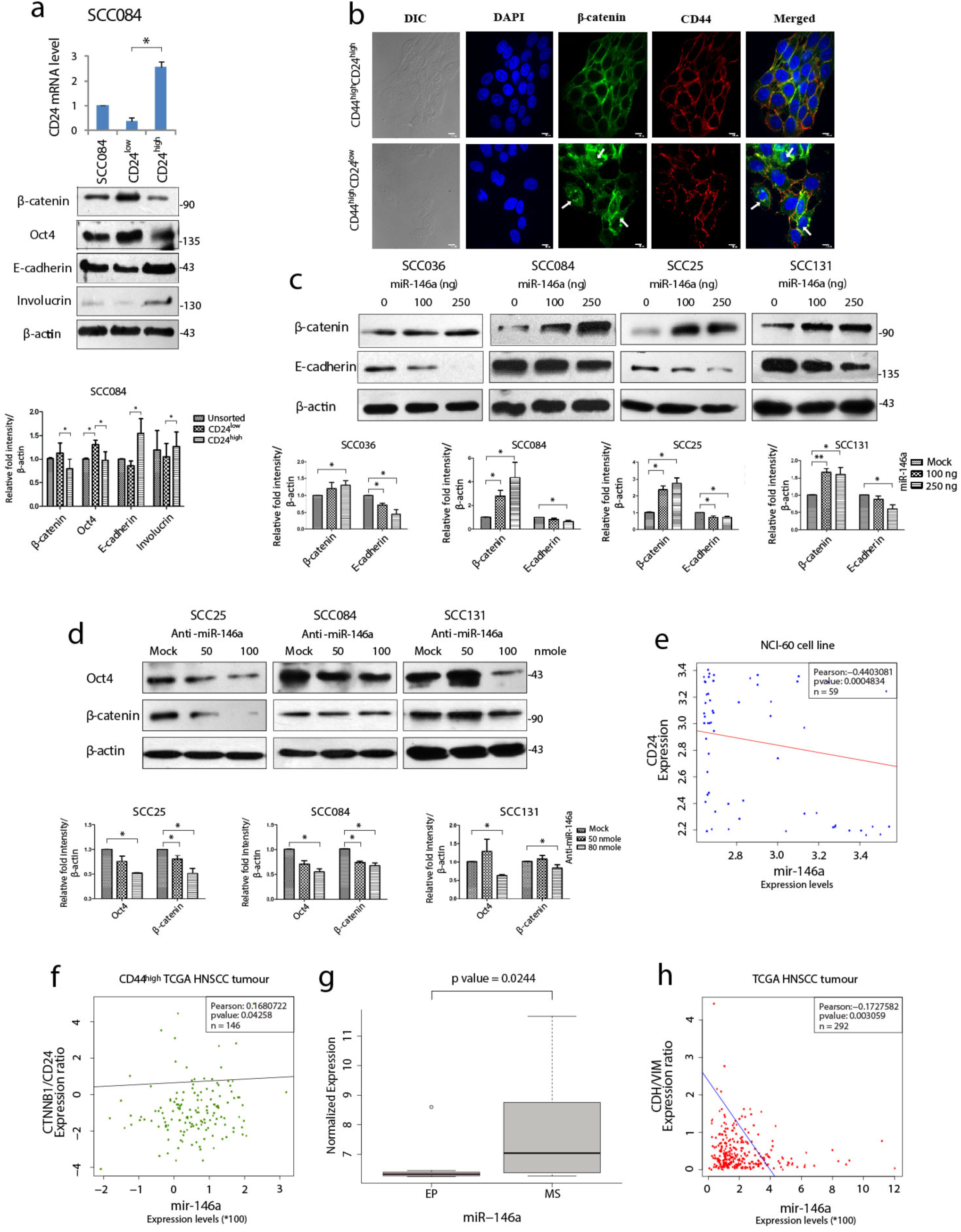
MiR-146a induced β-catenin/Wnt Signaling in CD44^high^CD24^low^ population. (**a**) Isolation of CD24^low^ and CD24^high^ cells from SCC084, as shown by qRT-PCR of CD24. Western blot images of β-catenin, Oct-4, E-cadherin and Involucrin in the respective populations along with its quantitative plot indicating the statistical data. (**b**) Representative confocal immunofluorescence images (60X magnification) of CD44^high^CD24^high^ and CD44^high^CD24^low^ subpopulation of SCC131 showing β-catenin (green) and CD44 (red) counterstained with DAPI (blue) (scale bar equal to 10 µm) (**c**) Each of the SCC cell lines were subjected to western blot analysis of β-catenin and E-cadherin upon increasing doses of miR-146a. (**d**) β-catenin and Oct-4 immunoblotting upon increasing doses of anti-miR-146a. Data normalized with β-actin. All the data has been graphically represented beneath the respective figures (* P<0.05, **P<0.01, *** P<0.001) (**e**) Association of miR-146a expression with CD24 in the NCI-60 cell lines (n=59) and (**f**) with β-catenin/CD24 ratio in the CD44^high^ tumors of the TCGA HNSCC patients (n=146). Statistical significance was determined by Pearson correlation test. Pearson correlation coefficient is shown in each plot. **(g)** Box plots showing miR-146a expression in NCI-60 cell lines classified as epithelial (EP) and mesenchymal (MS) subgroups. p value has been calculated using mann-whitney’s u-test. **(h)** Correlation (Pearson) of miR-146a with CDH1/VIM expression ratio based on the RNA-Seq data from 292 TCGA HNSCC specimens

The clue that β-catenin level might influence stemness in OSCC cells led us to investigate whether miR-146a regulates β-catenin. Indeed, over-expression of miR-146a lead to the dose dependent increment in β-catenin levels with concordant decrease in E-cadherin (Fig.3c). It is known that miR-146a targets the 3′UTR of Numb, a protein that promotes lysosomal degradation of β-catenin and Cleaved Notch1 [35].We indeed, confirmed reduced levels of Numb upon miR-146a over-expression along with stabilization of Cleaved Notch1(Figure S3e). Conversely, inhibition of miR-146a activity lead to β-catenin degradation along with the loss of Oct4 as expected (Fig.3d). Notably, we did not observe these changes upon mutant miR-146a over-expression (Figure S3f). To emphasize the contribution of β-catenin in miR-146a induced stemness, we transfected miR-146a in β-catenin shRNA expressing cells and found no change in expression of CSC markers (Figure S3g) as compared to that of non-silencing controls. Strikingly, the ability of anchorage independent growth induced by miR-146a was also dependent on the expression of β-catenin (Figure S3h) suggesting its tumorigenic role in OSCC.

To obtain clinical relationships among miR-146a, CD24 and β-catenin, we first checked the correlation between miR-146a and CD24 expression across the NCI-60 cell lines and found it to be significantly negatively correlated (Fig.3e). Moreover, examination of CD44^high^ HNSCC tumors from TCGA dataset revealed a positive correlation between miR-146a expression and β-catenin/CD24 ratio (Fig. 3f). Further, the observed down-regulation of E-cadherin upon miR-146a expression prompted us to address the miR-146a driven EMT phenomenon in OSCC. Indeed, miR-146a was found to be significantly over-expressed in the mesenchymal (MS) cell lines showing higher CD44 and lower CD24 expression[2], compared to the epithelial (EP) cell lines of the NCI-60 panel[26](Fig.3g). In addition, the miR-146a expression in TCGA tumor samples was negatively correlated with the E-cadherin to Vimentin ratio (Fig.3h). Based on these observations, we propose that miR-146a induced stemness and EMT in OSCC is mediated through lowering of CD24 followed by activation of β-catenin.

### 3.4 MiR-146a targets CD24 in oral CSCs

We considered CD24 as a putative target of miR-146a through which it might impart stemness in OSCC. Although, *in silico* identification of miRNA targets using the prediction software did not reveal CD24 as the probable target, we did find matching of miR-146a seed sequence in the CD24 3′UTR in miRANDA (Supplementary file 2). Despite one mismatch, the maximum free energy of miRNA-mRNA binding was favorable enough for hybridization and targeting (Fig. 4a). In Fig. 2c, we had already examined that CD24 expression was significantly depleted upon miR-146a transfection in a dose dependent manner in SCC036, SCC131 and SCC084 cells. Alongside, it was up-regulated upon inhibition of miR-146a in SCC131 cells (Fig.4b). We also observed remarkable changes in CD24 transcripts in both SCC131 and SCC084 upon modulation of miR-146a (Fig. 4c,d).To further confirm that CD24 is a direct target of miR-146a, we co-transfected luciferase reporter vector containing the 3′UTR fragment of CD24 gene with either miR-146a expressing vector or anti-miR-146a in SCC131 cells. As shown in Fig.4e, f and g, miR-146a over-expression reduced the luciferase activity of CD24 3′UTR, while miR-146a inhibitor elevated the same. On the contrary, transfection with mutated mir-146a did not alter the CD24 3′UTR luciferase activity significantly (Fig.4e, h). Thus, this data experimentally validates the ability of miR-146a to directly target CD24 gene by binding to its 3′UTR. This justifies the involvement of miR-146a in negative regulation of CD24 expression in oral cancer stem cells.

**Fig. 4.**
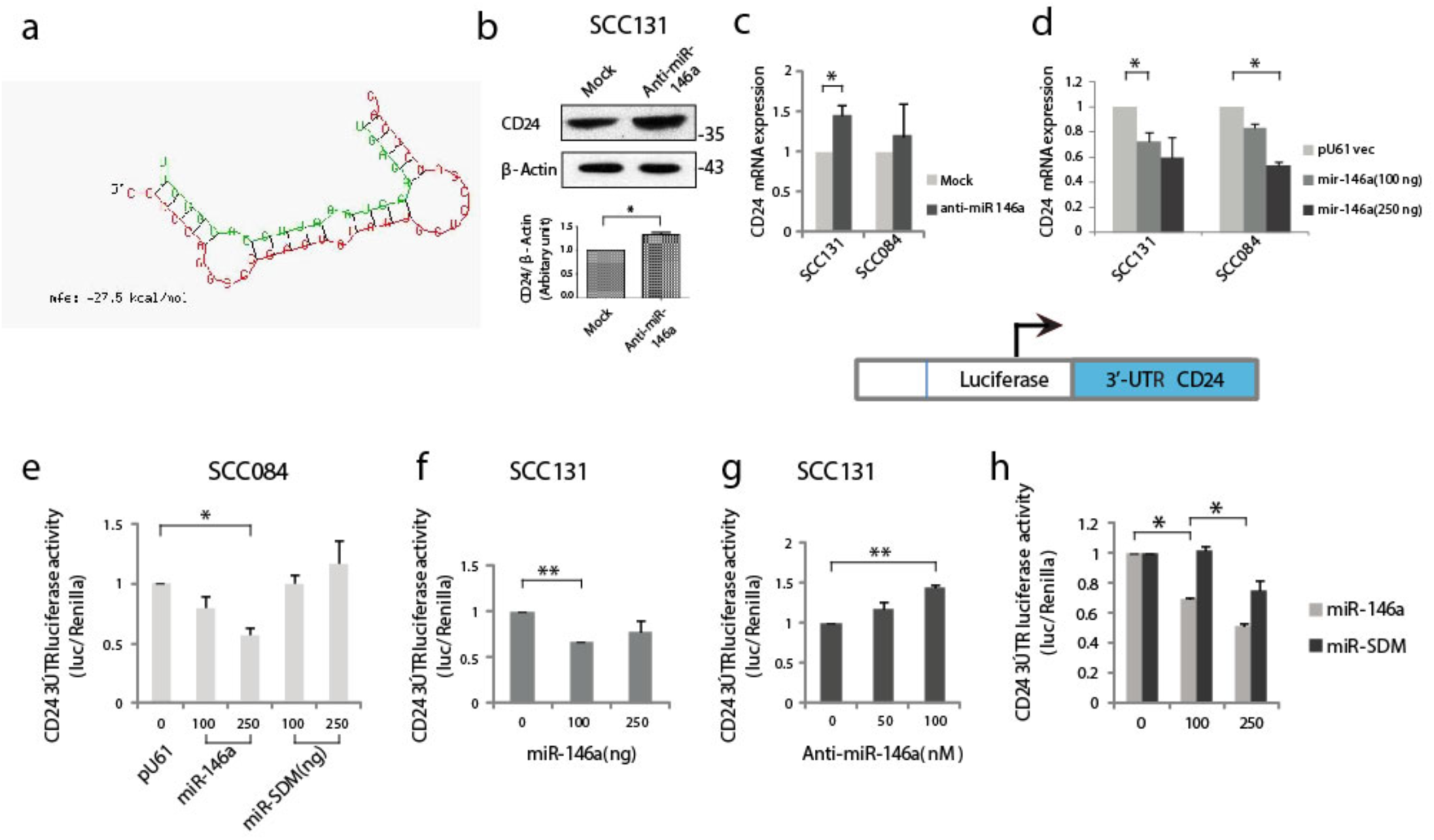
MiR-146a targets CD24 post-transcriptionally. **(a**) Secondary structure prediction of CD24 3′UTR base pairing with the seed sequences of miR-146a with negative free energy allowing favorable binding. (**b**) Immunoblotting of CD24 in SCC131 cells subjected to miR-146a knockdown showing significant up-regulation upon normalization with β-actin. (**c**) qRT-PCR to detect CD24 transcripts in the anti-miR-146a treated or (**d**) miR-146a over-expressed SCC131 and SCC084 cells.18srRNA served as an endogenous control and p-values calculated by Student’s t-test. Schematic representation of the CD24 3′UTR luciferase constructs (below). (**e**) Luciferase activity of the CD24 3′UTR reporter gene in SCC084 cells with increasing expression of miR-146a or miR-SDM (mutant miR-146a). (**f**) CD24 3′UTR reporter activity upon miR-146a over-expression and (**g**) miR-146a knockdown in SCC131 cells. Data in (**e**)-(**g**) are mean ±SD, normalized with pRL-TK vector and statistical significance was measured by paired Student’s t test (two-tailed). **(h)** Quantification of CD24 3′UTR luciferase activity in HEK293 T cells under miR-146a or miR-SDM over-expressed condition. Bar diagram represent the mean ± se calculated from three independent experiments. p-value was derived from Student’s t test.

### 3.5 Mir-146a stabilizes β-catenin by down-regulating CD24

To get a mechanistic insight into the miR-146a mediated β-catenin stabilization, CD24 was over-expressed in miR-146a over-expressing cells. Interestingly, CD24 not only abolished the stemness markers but also the expression of β-catenin and CD44 (Fig.5a and Figure S4a).This indicated that down-regulation of CD24 by miR-146a was instrumental in maintaining high β-catenin levels and consequently the downstream stemness phenotype. This was further exemplified as CD24 over-expression alone could lead to degradation of β-catenin protein and the associated stem cell markers, while the siRNA mediated knockdown of CD24 showed an opposite effect (Fig.5b,c). Wnt target genes such as *C-myc, CD44, CCND1* were detected at conspicuous levels upon CD24 knockdown, whereas significantly depleted upon CD24 over-expression in cells expressing miR-146a (Figure S4b, c). These observations suggest an inverse correlation between CD24 and β-catenin signalling in OSCC cells. It was further confirmed by the inhibition of miR-146a-induced β-catenin nuclear mobilization upon ectopic expression of CD24 (Figure S4d). In addition, miR-146a driven increased wnt reporter activity was found to be reduced upon CD24 over-expression (Fig. S4e). To further investigate its downstream effect on stemness phenotype, we performed an *in vitro* sphere-formation assay. We observed a considerably defective spheroid forming ability of miR-146a transfected cells in the presence of CD24 (Fig.S4f). Together, these observations suggest the possible contribution of CD24 in regulating Wnt pathway through β-catenin, thereby affecting CSC-like traits.

**Fig. 5.**
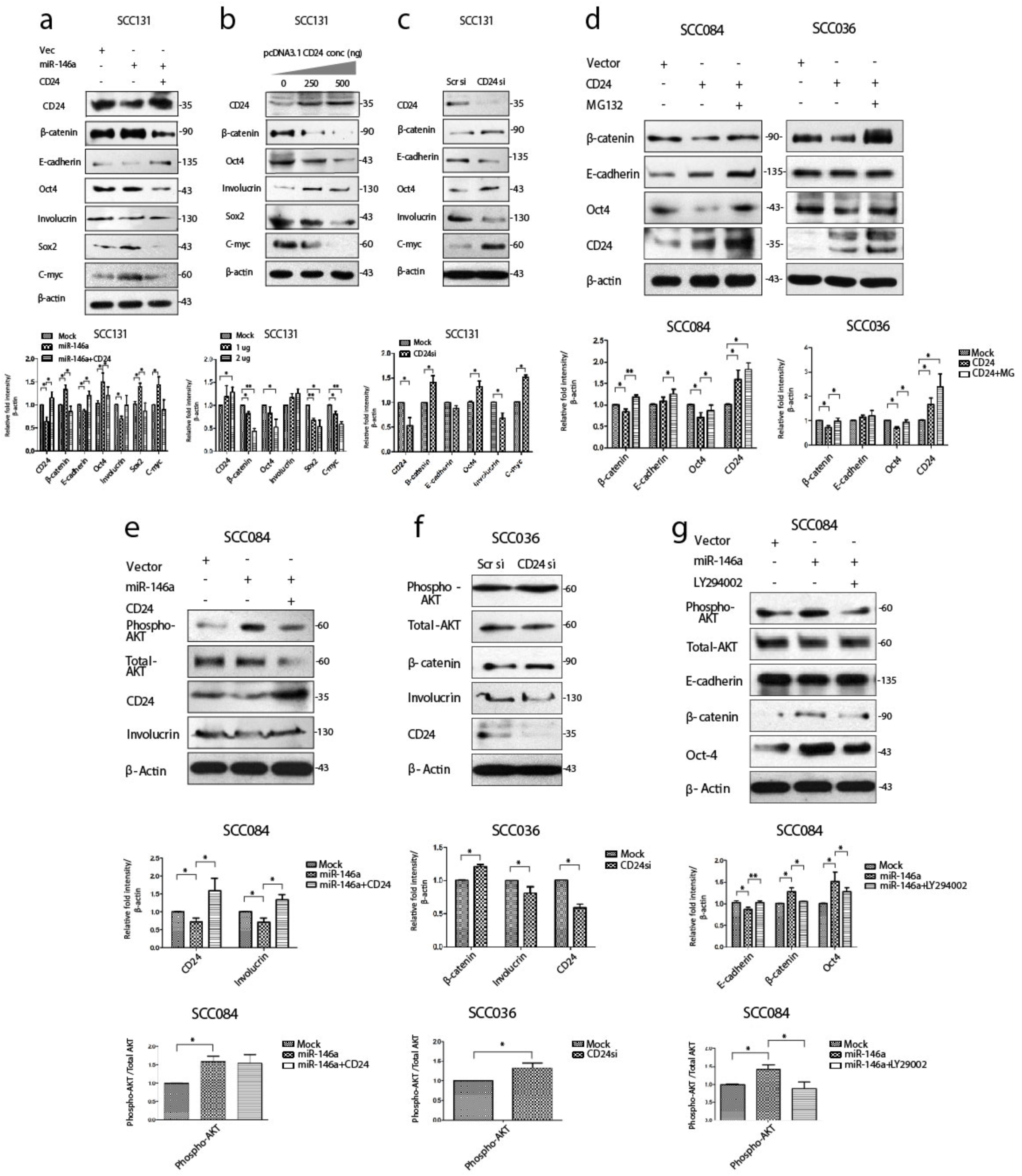
MiR-146a promotes stemness by down-regulating CD24 and leads to AKT mediated stabilization of β-catenin. (**a**) Immunoblot analysis of SCC131 transfected with either a vector control, miR-146a with or without CD24 showing rescued expression of β-catenin and E-cadherin as well as Involucrin Oct-4, Sox2, C-myc. (**b**) Immunoblotting of the same proteins with increasing dose of CD24 expression and (**c**) upon knockdown of CD24 in SCC131 cells. β-actin is loaded as an endogenous control. Histograms show fold changes in the densitometric values of band intensity and shown as means ± S.D. of 3 individual experiments. (**d**) Western blot analysis revealed degradation of β-catenin upon CD24 over-expression revived with MG132 treatment in SCC084, SCC036. (**e**) Phospho-AKT levels upon miR-146a alone or in combination with CD24. Total AKT is also shown. (**f**) Phospho-AKT and Total AKT levels upon siRNA mediated down-regulation of CD24 in SCC036 cells. (**g**) Effect of pAKT inhibitor upon β-catenin levels in SCC084 cells. Histograms showing fold change in the densitometric values of band intensity is represented as avg ± S.D. of n different experiments (n=3) (* P<0.05, **P<0.01).

### 3.6 Involvement of pAKT in CD24 mediated degradation of β-catenin

Next, to elucidate the cause of β-catenin reduction in the presence of CD24, we treated SCC084 and SCC036 cells with MG132, and found β-catenin levels returned to that of un-transfected controls (Fig.5d). This confirms that unlike Numb, CD24 degrades β-catenin in a proteasomally dependent manner. The restored β-catenin also re-established the expression of stem cell marker Oct4 irrespective of CD24 over-expression in both SCC036 and SCC084 cells (Fig.5d). E-cadherin levels, however, remained high in the presence of CD24, irrespective of β-catenin stability (Fig.5d). Notably, CD24 did not affect Numb expression, corroborating the independent participation of CD24 in regulating β-catenin (Figure S5a). To gain mechanistic insights into the CD24 mediated β-catenin degradation, we speculated that CD24 may lead to β-catenin destabilization via AKT-GSK-3β pathway[36, 37]. Towards this, we did find that CD24 over-expression rescued the miR-146a mediated increase in pAKT (Ser 473) activity (Fig.5e), although it remained stable during MG132 treatment (Figure S5b). On the contrary, knockdown of CD24 in SCC036 increased pAKT and β-catenin levels (Fig.5f). Moreover, pAKT was found to be accumulated in the CD44^high^CD24^low^ fraction of SCC25 cells thereby indicating its prior involvement in stemness (Figure S5c). In addition, we observed significant depletion of β-catenin upon treatment with pAKT inhibitor (LY294002) which again confirmed its regulation by miR-146a-CD24-pAKT loop (Fig.5g).

This was further supported by the soft agar tumorigenesis assays, which clearly showed that CD24 over-expression or AKT inhibition, both rescued the miR-146a induced colony formation in OSCC cells (Figure S5d). These cell line related observations was further validated using mouse xenograft model. To check the effect of miR-146a on *in-*vivo tumor formation, SCC084 cells harboring either an empty vector (SCC084/EV) or stably expressing miR-146a (SCC084/miR-146a), were generated and the over-expression of miR-146a with subsequent downregulation of CD24 was confirmed by both qRT-PCR and western blotting (Figure S6a, b and c). SCC084/EV and SCC084/miR-146a cells were then introduced in the right and left flanks of NOD/SCID mice respectively and allowed to form subcutaneous tumor (Figure S6d and e). A significant increase in tumor volume and tumor weight was observed in SCC084 xenografts stably over-expressing miR-146a suggesting enhanced stemness potential of these cells. (Fig 6a and b). Further, to explore the effect of AKT signaling on miR-146a induced tumor, mice with palpable tumors generated from SCC084/ EV and SCC084/miR-146a cells were treated with quercetin, a known PI3K/AKT signalling pathway blocker [38] (Fig S6d and e). As expected, the tumor formation ability of miR-146a cells was significantly attenuated *in vivo* upon administration of quercetin at regular intervals (Fig 6a and b). While there was no effect of quercetin on miR-146a and CD24 levels, β-catenin over-expression in miR-146a tumors was compromised in presence of quercetin (Figure S6f, g). In order to investigate the effect of CD24 upon the acquired stemness of miR-146a expressing cancer cells *in vivo*, we examined xenograft tumors generated from SCC084/miR-146a cells harboring CD24 expression construct (Figure S6a, b, c, h and i). Notably, compared to the control tumors, volume and weight of CD24 expressing SCC084/ miR-146a tumors were not significantly (p-value = 0.7360) altered (Fig 6c and d) suggesting loss of miR-146a driven stemness. Xenograft tumor subjected to qRT-PCR analysis confirms the overexpression of miR-146a and CD24 in these cells while β-catenin is down-regulated (Figure S6j and k). Collectively, these results indicate that CD24/AKT/β-catenin axis plays an important role in miR-146a regulated CSC-mediated tumor growth in-vivo.

**Fig. 6.**
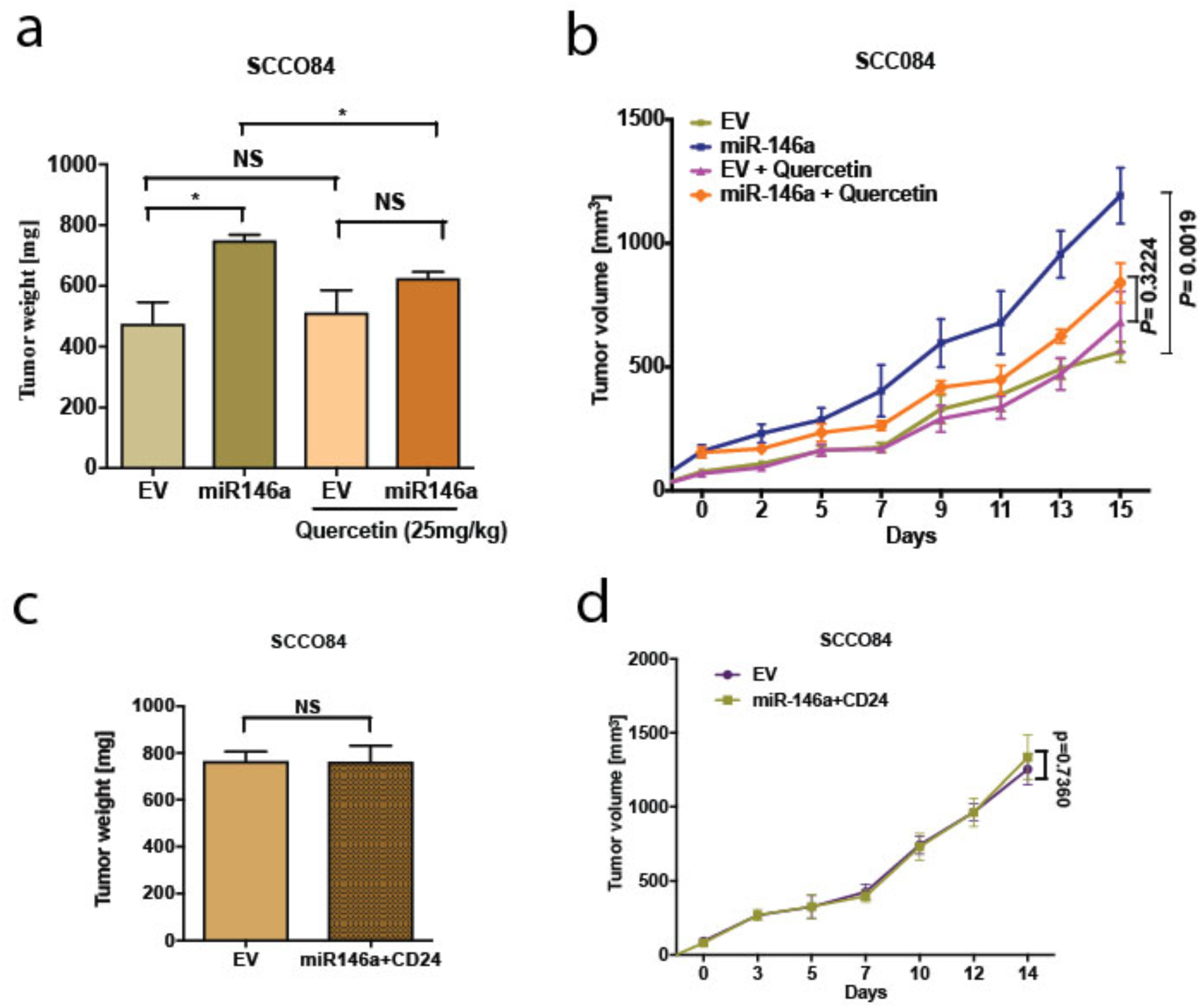
MiR-146a promotes *in vivo* tumor growth which is rescued upon CD24/AKT modulation (a) Bar graphs showing relative weight (mg) of the tumors xenograft tumors generated from control or miR-146a expressing SCC084 treated without or with Quercetin. Data represent mean ± SEM (n=4). P-values were computed using two-tailed Student’s t-test. (b) Line graph showing relative growth rate of tumors in response to Quercetin in SCC084 cells harboring either control vector or stably expressing miR-146a. Once tumors reached a palpable size, one set of mice were injected with Quercetin (10 mg/kg) intraperitoneally and after 10 successive treatment, change in tumor volume was measured at regular interval up to 20 days. Data represent mean ±SEM (n = 4) and *p*-values shown using Student’s t test. (c) Bar graphs showing relative weight (mg) of the tumors described in xenograft tumors generated from SCC084 harboring either control vector or stably expressing miR146a and CD24. Data represent mean ±SEM (n=4). P-values were assessed using 2-tailed Student’s t-test (d) Line graph showing relative growth rate of tumors described in (d). Data represent mean ±SEM (n = 4). For all the experiments, *p*-values were calculated in graph-pad prism5 using two-tailed Student’s t-test algorithm.

### 3.7 β-catenin transactivates miR-146a expression contributing to positive feedback loop

The upstream regulators of miRNAs have always been involved in feedback regulatory mechanisms and are not much investigated. Analysis of miR-146a promoter has revealed the binding sites for NF-κB, TCF4/β-catenin and STAT3, suggesting possible transcriptional modulation[18, 39, 40].β-catenin enhances stemness features by driving the intracellular levels of c-myc and other yamanaka factors (Fig.7a). This apparently contributes to the enhanced tumorigenic properties in response to high miR-146a levels. Interestingly, we found that β-catenin also promotes the expression of miR-146a, which might augment the stemness acquiring ability of the cancer cells (Fig.7a). However, expression of miR-146a was significantly reduced in the presence of both dnTCF4 and Numb which inhibits β-catenin binding to the promoter and degrades it respectively (Fig.7b). Change in miR-146a promoter activity under similar conditions suggests that β-catenin is involved in trans-activation of miR-146a promoter (Figure S7a). We hypothesize that β-catenin mediated induction of miR-146a contributes to β-catenin mediated CD24 reduction (Fig.7a). ChIP-qPCR assay confirmed that β-catenin binds to miR-146a promoter *in vivo* (Fig.7c). Further, the recruitment of β-catenin was found to be significantly enhanced in absence of CD24 (Fig.7c). Moreover, miR-146a promoter activity was significantly increased in presence of miR-146a, while reduced upon ectopic expression of CD24 (Fig. 7d). However, the constructs with either mutated or deleted TCF4 binding sites showed no significant difference in promoter activity (Figure S7b). This may be due to the alternating levels of β-catenin which shoots up in miR-146a over-expression condition and gets depleted in presence of CD24. Transient ChIP assays with the same luciferase constructs also confirmed that β-catenin indeed binds to miR-146a promoter, which is impeded upon CD24 over-expression (Fig. 7e). These data positively confirm the feedback activation loop by β-catenin that further trans-activates miR-146a expression to shift the equilibrium towards CSC maintenance.

**Fig. 7.**
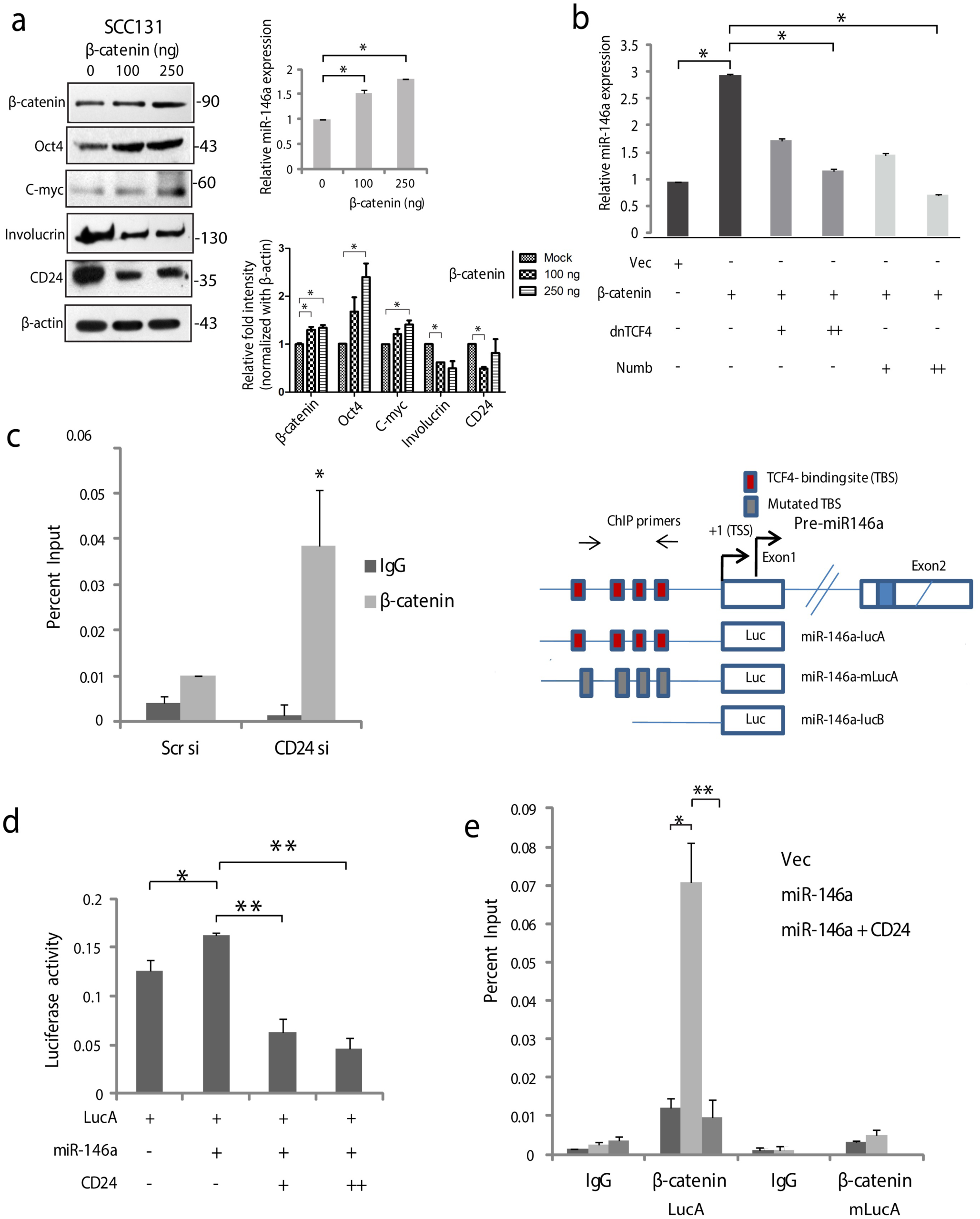
β-catenin transactivates miR-146a mediating a positive feedback loop. (**a**) Increase in stemness markers upon β-catenin over-expression as revealed by western blot and the densitometric analysis of its band intensities (right below). Data was normalized with corresponding β-actin. Concomitant miR-146a expression in β-catenin transfected SCC131 as quantified by qPCR. (**b**) Increase in miR-146a transcripts upon β-catenin over-expression is dose dependently inhibited in the presence of either dnTCF4 or Numb. (**c**) Chromatin Immunoprecipitation assays in SCC084 cells transfected with either scramble siRNA or CD24 siRNA showing recruitment of β-catenin upon endogenous miR-146a promoter. Schematic of miR-146a promoter locus and reporter constructs namely LucA: wild-type; mLucA: TCF-4 binding site (TBS) mutation; LucB: TBS deletion[18]. (**d**) Relative luciferase activity of LucA in SCC084 cells under various transfections as indicated. Data represent average of n=3 individual experiments (along with technical replicates). (**e**) Transient ChIP assay with same constructs as shown. The percentage enrichment of amplified product was normalized to input and graphically presented. Data represent mean ± sd and n=3 different experiments (along with technical replicates\**P*< 0.05, \*\**P*< 0.01). We used Student’s *t*-test calculate *p-*value.

## 4 Discussion

Oral cancer progression has been largely attributed to both genetic and epigenetic heterogeneity. Tumorigenic cells can arise from the non-tumorigenic cancer cells owing to spontaneous conversion to a stem-like state[41]. The origin and plasticity of such cells, called cancer stem cells (CSCs), have always been a matter of debate. Nevertheless, CSCs are known to be responsible for chemo-resistance, tumor recurrence and metastasis. Detail molecular characterization of CSCs is therefore of paramount importance for eliminating them from its roots. CD24 has been routinely used with CD44 for the prospective isolation of CSCs in colorectal, prostate and breast cancers [42]. Given the critical role of cellular miRNAs in regulating CSC characteristics, current anticancer therapies have provided an important avenue towards exploiting them for effective cellular targeting[43]. Targeting of CD44 by miRNAs in NSCLC, prostate and ovarian cancer has been previously demonstrated to attenuate stemness[14, 44, 45]. However, miRNA mediated regulation of CD24 remains to be determined.

Consistent with its oncogenic functions, miR-146a promotes symmetric division of colorectal CSCs, thereby promoting stemness [18]. The miRNA is also involved in development of melanoma by activating Notch1 signaling leading to drug resistance [46]. However, little is known about its role in regulating expression of CSC-related CD markers in oral cancer. In the present study, we detected significantly higher expression of miR-146a in CD44^high^CD24^low^ population of OSCC cell lines as well as in tumor specimens. We therefore investigated whether miR-146a expression maintains CSC traits or miR-146a accumulation is a consequence of induced stemness. Notably, ectopic expression of miR-146a induce CSC-phenotype i.e. CD44^high^CD24^low^ population together with increased β-catenin activity in OSCC cell lines. CD44 is a well-known transcriptional target of β-catenin along with C-myc and CCND1[18], which clearly indicates a molecular link with miR-146a induced CD44 expression. However, the effect of miR-146a upon CD24 expression under these conditions was particularly intriguing. Hence, we examined whether CD24 is a direct target of miR-146a and experimentally confirmed that miR-146a binds to the 3′UTR of CD24 thereby repressing it post-transcriptionally. Loss of E-cadherin upon miR-146a over-expression and positive correlation with the mesenchymal marker vimentin was also evident. Hence, in addition to its novel role in acquiring stemness, our results re-confirmed miR-146a as a key regulator of EMT[47].

Wnt/β-catenin has shown great potential for CSC-targeting in cancer[34]. Our study shows that CSC characteristics in OSCC is attributed to the elevated β-catenin along with depleted CD24. The anticipation that CD24 leads to proteasomal degradation of β-catenin was found to be true and apparently it also abolished the β-catenin mediated stemness. This is a novel functional interaction through which miR-146a regulates β-catenin in oral cancer cells. Our study, thus points towards the tumor-suppressor functions of CD24, supporting our previous observation of reduced CD24 expression in oral tumors compared to the normal tissue (our own data) [2]. Although growth inhibition was achieved by knocking down CD24 in colorectal and pancreatic cancer[48], no such effects were observed in oral cancer. Perhaps the variable cell-type specific distribution pattern underlies the paradoxical role of CD24 in oral cancer[49]. Activated PI3K-AKT pathway is one of the primary events in carcinogenesis[50]. Its contribution to stem cell self-renewal and proliferation has also been extensively studied[51]. Further, signaling pathways like WNT are often linked with AKT activation that eventually contribute to expression of stem cells-related factors, chemo-resistance genes, and CSC markers[50, 52]. Here we show that CD24, the cell surface CSC marker lie upstream of AKT protein, similar to that of TWIST and FOXO transcription factors which is also known to inhibit CD24[36, 53–55]. However, the precise mechanism by which expression/stability of AKT protein is regulated by CD24 is unknown.CD24 has been shown to possibly modulate phospho-AKT levels [56], which might affect its downstream targets such as GSK-3β[37]. Activated GSK-3β mediates phosphorylation and ubiquitination of β-catenin, thereby leading to its degradation[37]. Therefore, it was incumbent on us to ask whether CD24 induce AKT and subsequently affect β-catenin stability in miR-146a induced oral CSCs. Indeed, MG132 treatment was found to re-stabilize β-catenin by relieving pAKT inhibition in cells over-expressing CD24. Moreover, direct AKT inhibition in the miR-146a transfected cells depleted β-catenin, irrespective of CD24 level suggesting AKT is downstream of CD24. Thus, we logically elucidated the molecular mechanism underlying CD24 mediated β-catenin degradation in oral cancer cells. We have specifically shown that CD24 over-expression decrease levels of phospho-AKT leading to β-catenin instability. The role of miR-146a/CD24/AKT/β-catenin axis in maintaining the oral cancer stem cell populations is thus mechanistically evident. Studies from in-vivo tumor model system also confirms that these molecular mechanisms directly affect tumorigenesis.

Further, the recruitment of β-catenin onto miR-146a promoter was found to be negatively regulated by CD24 which might contribute to the fine tuning of stemness. These results clearly establish a cross-regulatory network between miR-146a and β-catenin, governed by a stem-related marker, CD24 in OSCC cells. Our study thus provides strong evidences which suggest that miR-146a promotes CSC characteristics of oral cancer cells by down-regulating CD24. Repression of CD24 leads to AKT stabilization followed by activation of Wnt/β-catenin signaling. Based on our observation, we propose a model wherein, AKT activity is an important determinant of miR-146a dependent β-catenin signaling (Fig.8). It should be noted, however, that β-catenin mediated CSC induction might be due to the induction of miR-146a expression or vice-versa. Taken together, the present study highlights a novel mechanism of miR-146a mediated self-renewal capacity of Oral CSCs that may have a prognostic or therapeutic value in oral cancer.

**Fig. 8.**
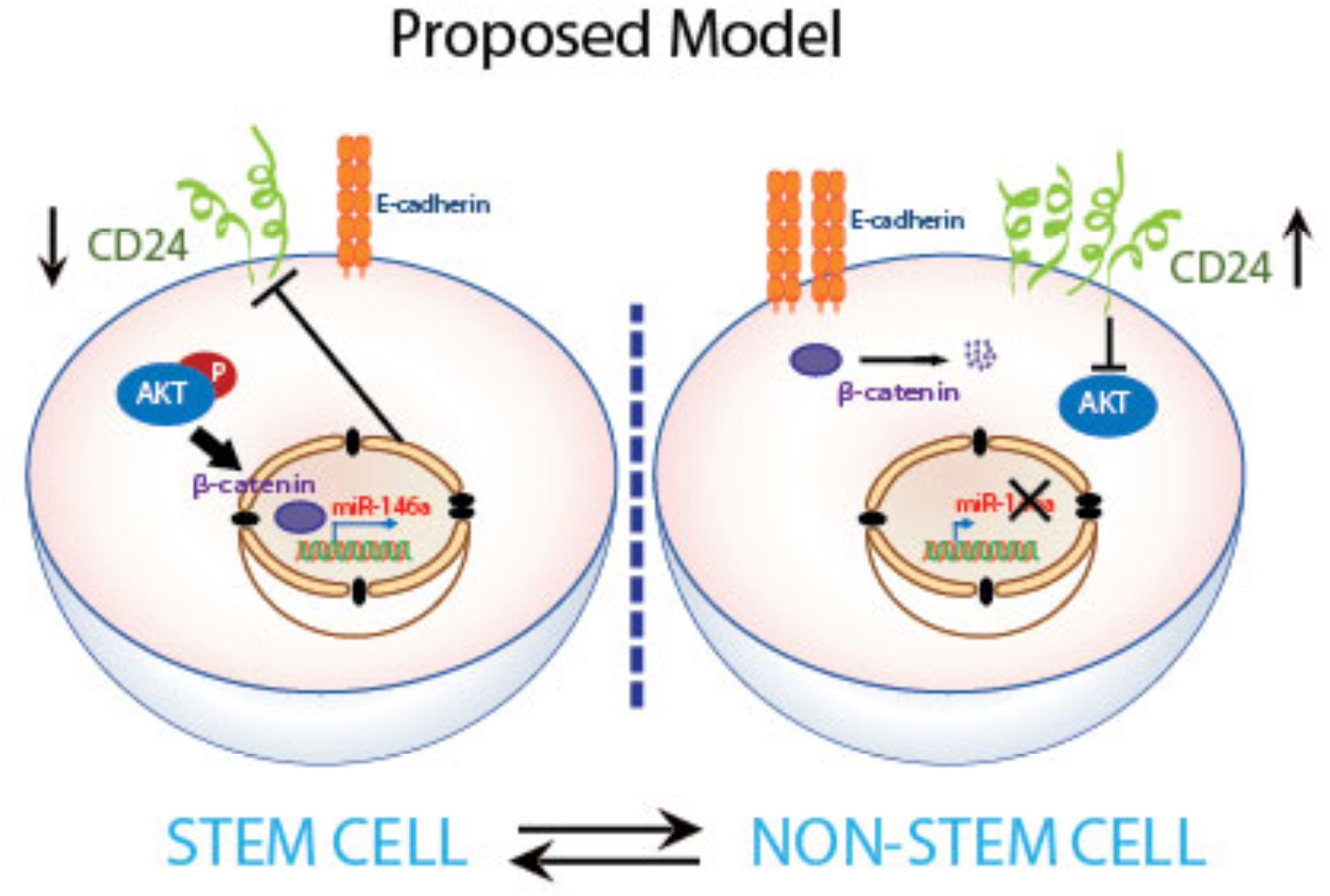
Model–schematic representation of transient inter-conversions between stem and non-stem oral cancer cells. Cancer stem cell induction may be triggered with over-expression of miR-146a that targets CD24 and leads to β-catenin protein stabilization via AKT activation. Wnt signalling intermediates promote stem-like de-differentiated state in the tumor cells. Accumulation of β-catenin might further drive miR-146a expression and amplify the stemness characteristics. Stochastic intra-cellular signals may induce differentiation, probably by aberrant activation of CD24. This leads to decline in pAKT levels and hence proteasomal degradation of β-catenin with subsequent loss of miR-146a.

## Supplementary information

**Supplementary File 1**: List of Primers for the qRT-PCR (Excel File)

**Supplementary File 2**: Supplementary Figure and Figure Legends (S1, S2, S3, S4, S5, S6, S7)

**Supplementary File 3**: Analysis of TCGA data for Head and Neck Squamous Cell Carcinoma patients (Excel File)

**Supplementary File 4**: Snapshots of miRANDA analysis for miR-146a-5p seed sequence and CD24 3′UTR matching showing their free energy of binding (pdf)

**Supplementary File 5**: Supplementary methods

## Author contributions

SG and SR conceived and designed the study. Some experiments were designed by DG and SD. Experiments, data collection and statistical analyses were performed by SG, DG, SD and PD. Some experiments were performed by RB and MG. The manuscript was written and edited by SG, DG, SD, GC and SR. All authors read and approved the final manuscript.

## Acknowledgements

We thank Prof. Nitai. P. Bhattacharjee (SINP), Dr. Muh-Hwa Yang (National Yang-Ming University, Taiwan) and Dr. Heike Allgayer (University of Heidelberg, Germany) for providing us the necessary plasmids. We thank Dr. Raghunath Chatterjee (Indian Statistical Institute, Kolkata, India) for performing the RNA-hybrid analysis in miRANDA software. We thank Tanmoy Dalui and Diptadeep Sarkar (CSIR-Indian Institute of Chemical Biology, Kolkata, India) for helping us with Flow Cytometry and Confocal Microscopy respectively. We thank Dr. Arindam Datta (CSIR-Indian Institute of Chemical Biology, Kolkata, India) for assisting TCGA and NCI-60 dataset analysis. The work was supported by CSIR—Mayo Clinic Collaboration for Innovation and Translational Research Grant CMPP-08 and J.C. Bose National Fellowship grant JCB/2017/000005 awarded to S. Roychoudhury. S.G, D.G and P.D. are supported by fellowship from the Council of Scientific and Industrial Research (New Delhi, India). Mouse experiments and SD, RB and MG are supported by CSIR-fellowship in NCCS, Pune.

