## Supplementary File 2 for "MiR-146a-dependent regulation of CD24/AKT/β-catenin axis drives cancer stem cell phenotype in oral squamous cell carcinoma"

##### Supplementary Figures and Figure Legends

**Supplementary Figure S1**

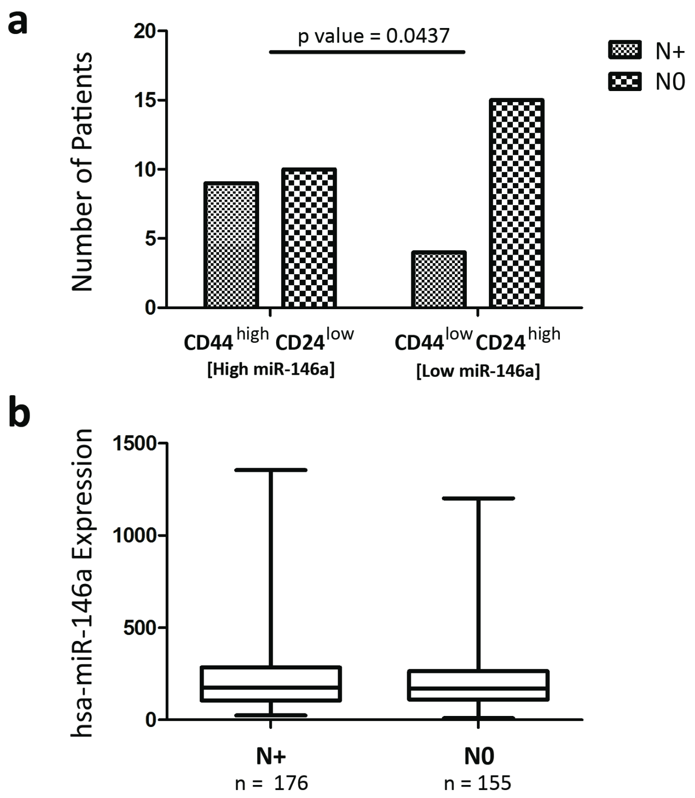


**Figure S1.** (**a**) Bar graph showing distribution of patients with node positive (N+) and node negative (N0) status within the $CD44^{high}CD24^{low}$ and $CD44^{low}CD24^{high}$ group as classified in Figure. 1c. p-value was calculated using chi-square test. **(b**) Box plots showing the expression of miR-146a across the node positive (N+) and node negative (N0) patients from the TCGA HNSCC ("n" represent no. of patients in each group).

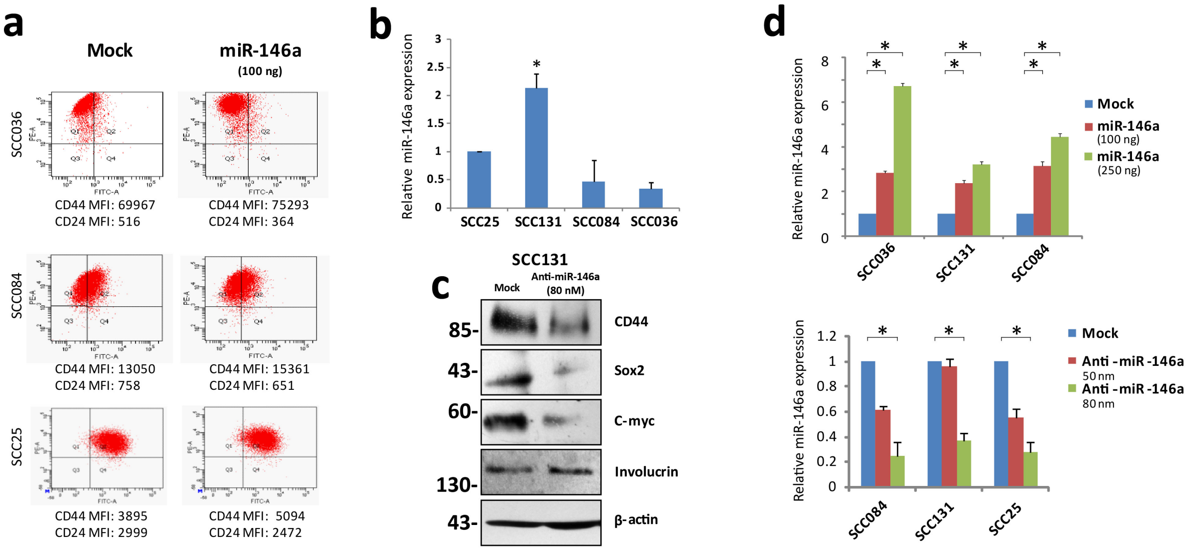


**Figure S2.** (**a**) Representative flow cytometry profile of SCC25, SCC084 and SCC036 cell lines with or without miR-146a over-expression stained with CD44 (PE) and CD24 (FITC). Mean fluorescence values are indicated for each cell line. (**b**) Endogenous levels of miR-146a were quantified in all the SCC cell lines by qRT-PCR. (**c**) Representative Western blot images of CD44, Sox2, C-myc, Involucrin in UPCI: SCC131 transfected with anti-miR-146a. (d) MiRNA specific cDNA derived from SCC131, SCC084, and SCC25 upon miR-146a over-expression or knockdown were subjected to qRT-PCR to confirm changes in experimental miR-146a levels. Relative expression values were normalized to those of U6snRNA. Data shown is average ± sd for three independent experiments. Student's t test was used to compute p-values (*$P$<0.05, **$P$<0.01 and ***$P$<0.001).

### Supplementary Figure S3

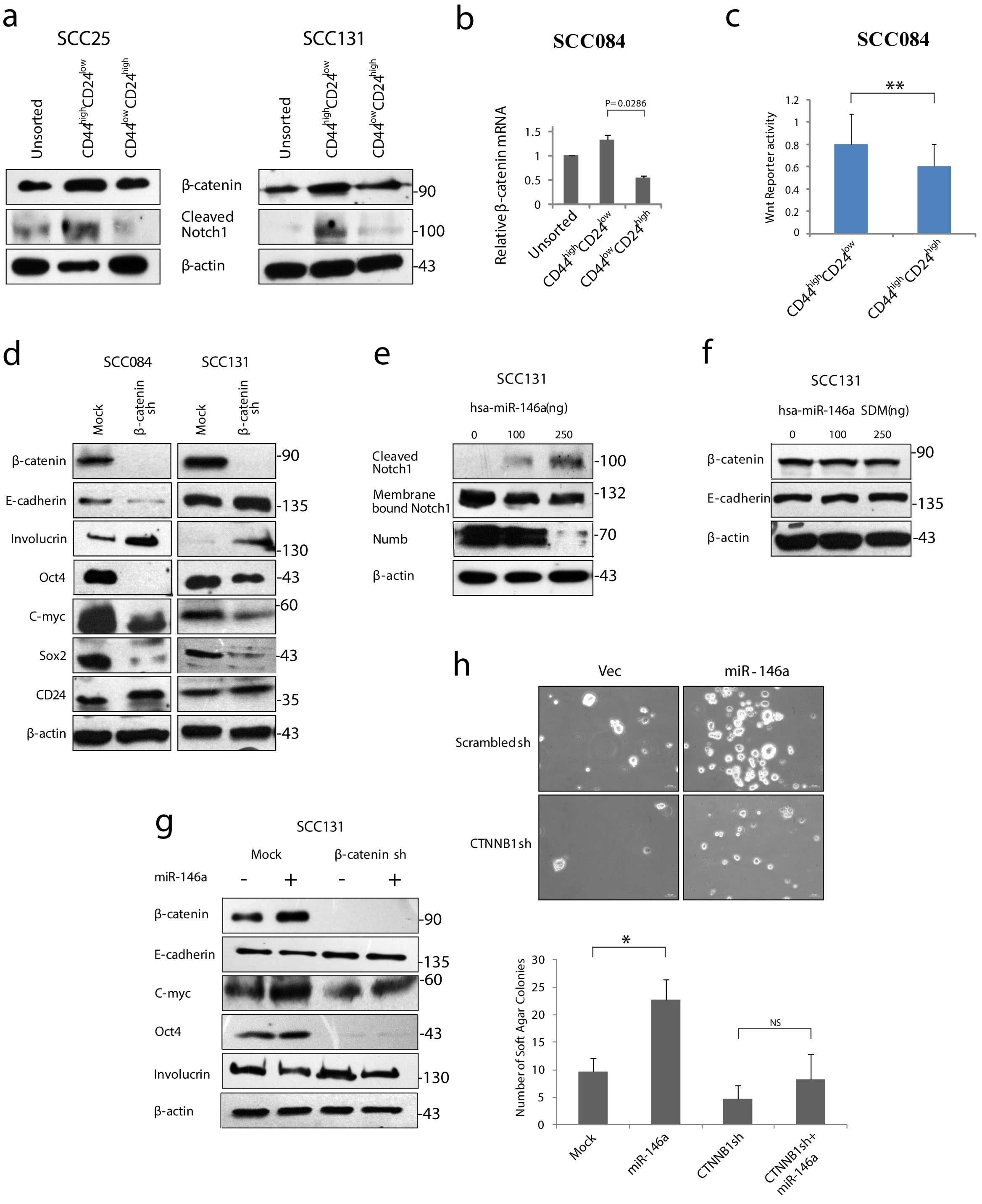

**Figure S3.** (**a**) Whole cell lysates from CD44$^{high}$CD24$^{low}$ and CD44$^{low}$CD24$^{high}$ population of SCC25 and SCC131 were subjected to western blot of Cleaved Notch1 and β-catenin and intensity normalized to β-actin. (**b**) Increased expression of β-catenin in the CD44$^{high}$CD24$^{low}$ compared to the CD44$^{low}$CD24$^{high}$population of SCC084 as measured by qRT-PCR. Data is represented as mean ±sd of at least three independent experiments. (**c**) Relative TOP/FOP luciferase activity in the CD44$^{high}$CD24$^{low}$ and CD44$^{high}$CD24$^{high}$ populations. The *P*- value was calculated using Student's *t*-test. (**d**) β-catenin knock-down cells (SCC131 and SCC084 β-catenin sh) along with control (SCC131 and SCC084 scrambled shRNA) cells were lysed and subjected to immunoblotting with stem cell markers. Silencing of β-catenin was also ascertained and β-Actin was used as a loading control. (**e**) Western blots of Cleaved Notch1, membrane bound Notch1 and Numb in miR-146a over-expressing SCC131 cells as a transfection control. (**f**) Immunoblotting of β-catenin and E-cadherin upon miR-146a-SDM transfection. (**g**) Effect of miR-146a transfection on the stemness markers in SCC131 (non-silencing and β-catenin sh) cells by Western blotting. The immunoblot experiment presented is one representative out of two independent experiments. (**h**) Soft agar colonies formed by scrambled and β-catenin shRNA transduced SCC131 with or without miR-146a transfection, followed by their quantification (p value calculated by Student's t test). Scale bar=50 µm.

**Supplementary Figure S4**

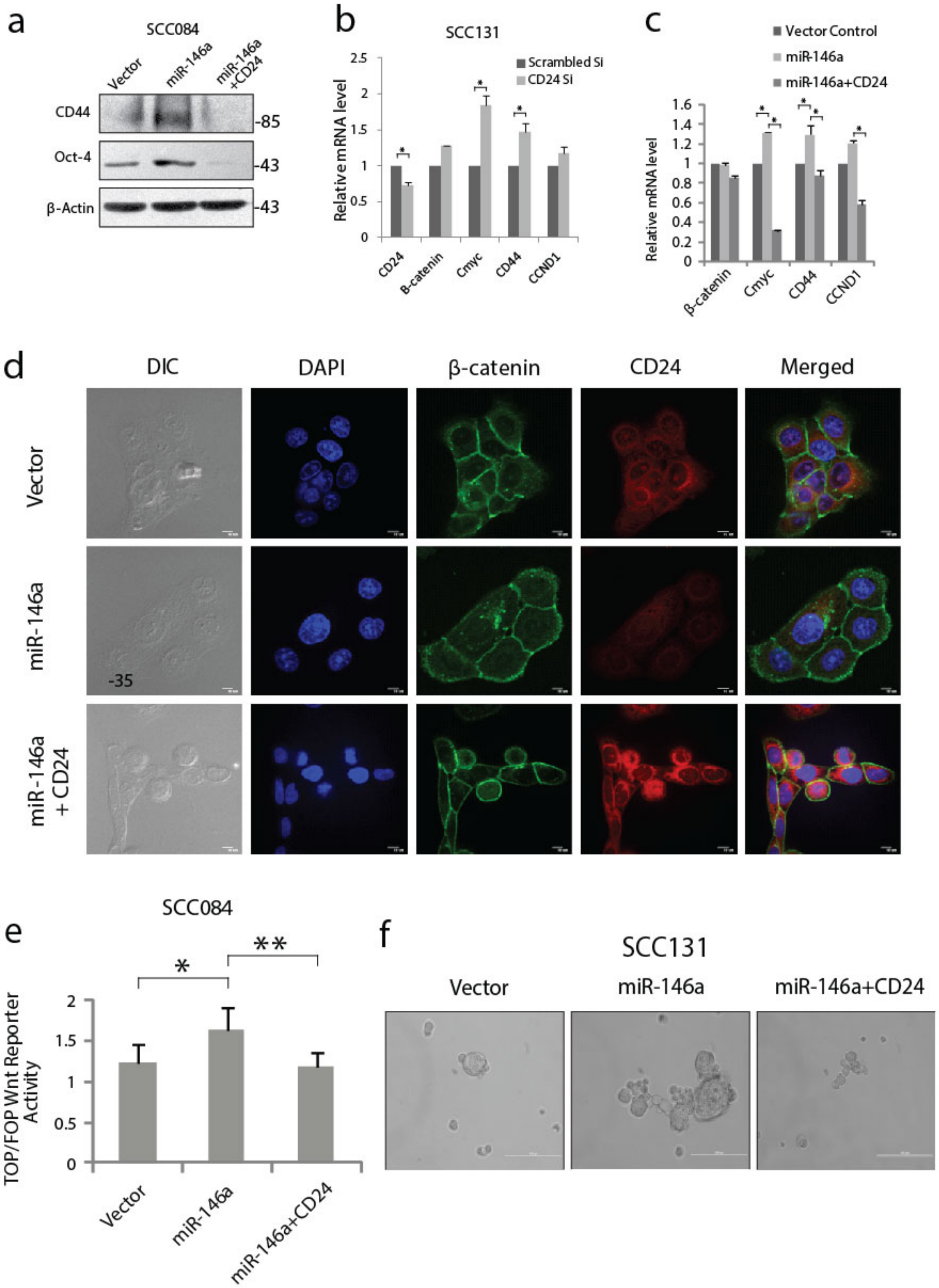

**Figure S4.** (**a**) Effect of CD24 on CD44 and Oct4 levels in the presence of miR-146a in SCC084 cells.(**b**) Total RNA extracted from SCC131 transfected with either non-target siRNA or CD24 siRNA were subjected to qRT-PCR of β-catenin along with its targets. **(c)** Quantification of β-catenin and Wnt target gene (C-myc, CD44 and CCND1) by qRT-PCR transfected with control vector, mir-146a alone or in combination with CD24. Data represent mean ± se, n=2 independent experiments. Statistical significance was calculated by Student's t test. (**d)** Representative images of immuno-fluorescent staining of β-catenin and CD24 in SCC131 transfected with a control vector or miR-146a and miR-146a co-expressing CD24 (Red, CD24; green, β-catenin; blue DNA; Scale bars, 10 μm). (**e**) Wnt reporter activity as measured by Top-Flash vs Fop-Flash luciferase construct in SCC084 under different transfections as indicated. (**f**) Sphere formation ability under similar conditions as described in (e) (Scale bar=200 μm).

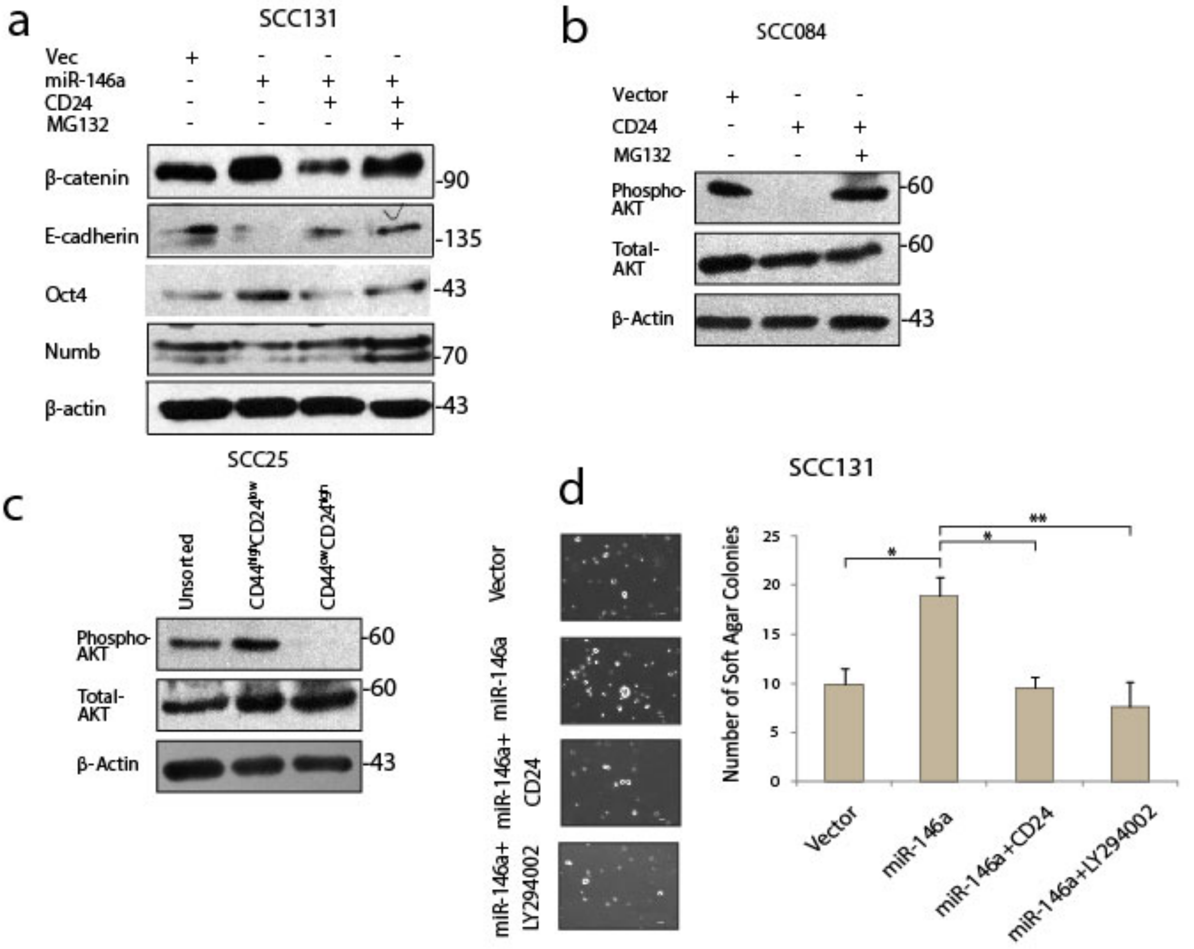


**Figure S5.** (**a**) Effect of CD24 upon Numb with or without MG132 in SCC131 cells. (**b**) Effect of CD24 upon Phospho-AKT and Total AKT levels in SCC084 cells with or without MG132. (**c**) Phospho-AKT levels in the CD44$^{high}$CD24$^{low}$ and the CD44$^{low}$CD24$^{high}$ subpopulation of SCC25. **(d)** Representative pictures of the colonies from the soft agar assay performed under given transfections and treatment conditions as shown (Left) (Scale bar=100 µm). Number of similar sized colonies were counted and avg ± sd were plotted and graphically represented (* P<0.05)

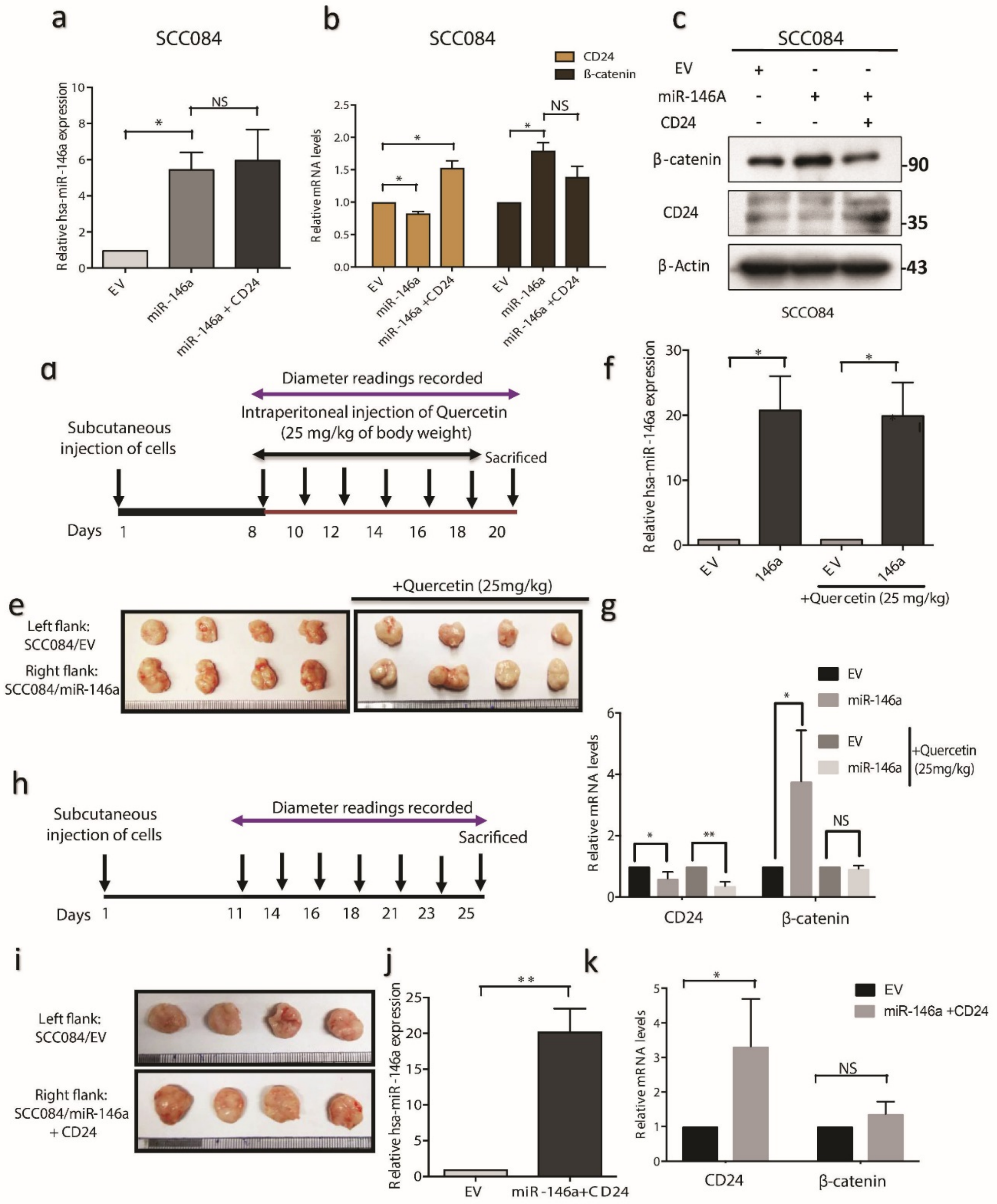
a
SCC084
Relative hsa-miR-146a expression
EV
miR-146a
miR-146a + CD24
NS
b
SCC084
CD24
β-catenin
Relative mRNA levels
NS
c
SCC084
EV
miR-146A
CD24
β-catenin
CD24
β-Actin
90
35
43
SCCO84
d
Diameter readings recorded
Intraperitoneal injection of Quercetin
(25 mg/kg of body weight)
Subcutaneous
injection of cells
Sacrificed
Days 1 8 10 12 14 16 18 20
f
Relative hsa-miR-146a expression
EV
146a
+Quercetin (25 mg/kg)
e
+Quercetin (25mg/kg)
Left flank:
SCC084/EV
Right flank:
SCC084/miR-146a
g
EV
miR-146a
+Quercetin
(25mg/kg)
Relative mRNA levels
CD24
β-catenin
h
Subcutaneous
injection of cells
Diameter readings recorded
Sacrificed
Days 1 11 14 16 18 21 23 25
i
Left flank:
SCC084/EV
Right flank:
SCC084/miR-146a
+ CD24
j
Relative hsa-miR-146a expression
EV
miR-146a+CD24
k
Relative mRNA levels
EV
miR-146a +CD24
CD24
β-catenin
NS

**Figure S6.** (a) QRT-PCR showing relative expression levels of miR-146a in cells stably expressing either miR-146a alone or miR-146a and CD24.(b) Relative mRNA expression levels of CD24 and β-catenin in the same conditions as in (a). (c) Immunoblots showing β-catenin and CD24 levels in SCC084 cells as described in (a). (d) Schematic of subcutaneous injection of control or miR-146a expressing cells in the dorsal flanks of mice and timeline of quercetin treatment/recordings of tumor volume. (e) Representative images showing tumor obtained from NOD/SCID mice inoculated with either SCC084/EV or SCC084/146a without or with Quercetin treatment. (f) QRT-PCR data showing relative expression levels of miR-146a in SCC084/EV and SCC084/146a tumors and in response to Quercetin. (g) QRT-PCR data showing relative expression levels of β-catenin and CD24 in SCC084/EV and SCC084/146a tumors in conditions as described in (f). (h) Scheme showing control or miR-146a and CD24 co-expressing cells and various times of tumor measurements. (i) Representative images of tumors generated from SCC084 cells either harboring control empty vector or stably expressing miR-146a and CD24. (j, k) QRT-PCR showing relative expression levels of miR-146a (j), CD24 and β-catenin (k) in tumors as described in (h). β-actin served as the loading control. Bar graphs represent mean SD; n = 3; two-tailed Student's t-test: *$P < 0.05$, **$P < 0.01$, ***$P < 0.001$, NS, non-significant.

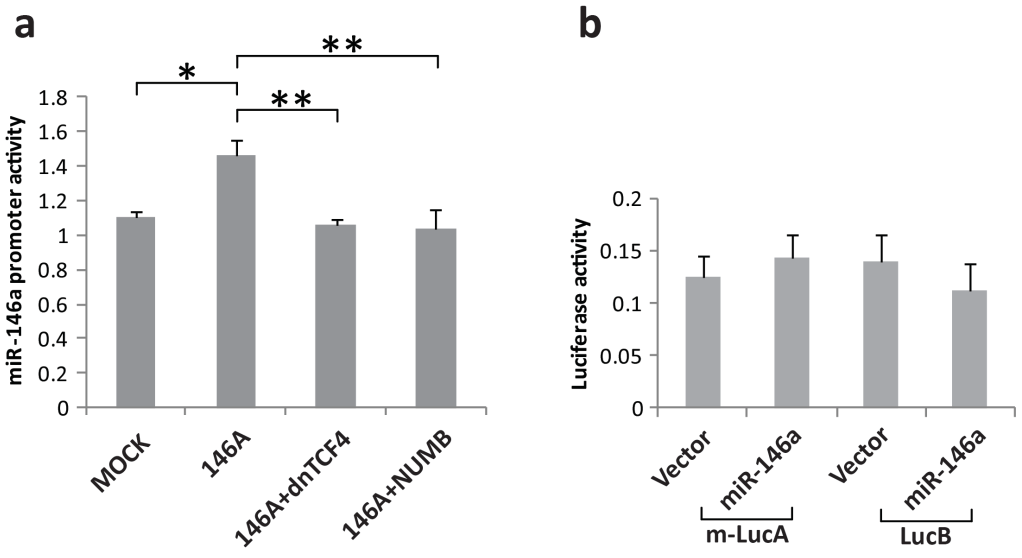


**Figure S7.** (**a**) LucA activity in the miR-146a over-expressing cells diminished upon dnTCF4 or Numb transfection. (**b**) m-LucA and LucB activity with or without miR-146a in SCC084 cells and relative luciferase activity measured. Data represent mean ± sd, n=2 independent experiments (each experiment contains 3 technical replicates). p value determined by student's t test.
