## Supplementary File 5 for "MiR-146a-dependent regulation of CD24/AKT/β-catenin axis drives cancer stem cell phenotype in oral squamous cell carcinoma"

**Supplementary Material and Methods:**

**Magnetic Assisted Cell Sorting**

MACS were performed as described previously (Ghuwalewala et al., 2016) to isolate the CD44^high^CD24^low^ (stem) and CD44^low^CD24^high^ (non-stem) populations in various oral cancer cell lines.

**Generation of Stable Knock-down cells**

Third generation lentiviral packaging vector, pLKO.1 puro short hairpin RNA (shRNA) for human beta-catenin and non-silencing control were kind gifts from Dr.Mrinal Kanti Ghosh, IICB. Lentivirus particles containing assembled CTNNB1 shRNA were generated in Hek293T cells using Tran-lentiviral packaging kits (Dharmacon, Thermo Scientific) as per manufacturer’s protocol. SCC131, SCC084 and SCC25 cells were then infected with viral supernatant using Polybrene (sigma) at 8 µg/ml. The cells were maintained under puromycin (1 µg/ml) (Gibco) selection. Stable selected clones were subsequently verified for loss of β-catenin expression by Western blotting and then used for further experiments.

**Soft agar assay**

Both the β-catenin and Non-silencing shRNA controls were independently transfected with miR-146a expressing vector and empty vector control respectively. SCC131 cells were transfected with empty vector control, miR-146a expressing vector and co-transfected with CD24 respectively. For AKT inhibition, miR-146a transfected cells were treated with LY294002, 4 hours prior to seeding. Twenty thousand cells from each set were counted and seeded in the 0.4% upper agarose medium layered on the top of 0.8% lower agarose medium in six-well plates. The above layer was kept moist with 1X medium at 37 ºC and colonies were pictured after 15-21 days under microscope (Leica CTR4000) at 20x magnification. The colonies were counted from random fields of three independent experiments and avg ± sd was plotted and statistically analysed.

**Composition of ChIP buffers:**

SDS lysis buffer (1% SDS,10 mM EDTA, 50 mM TRIS, pH 8.1)

ChIP dilution buffer (0.01% SDS, 1.1% TRITON X-100, 1.2 mM EDTA, 16.7 mM Tris-HCl,pH 8.1, 167mMNaCl)

Cold low salt immune complex wash buffer (0.1% SDS, 1% Triton X-100, 2 mM EDTA, 20 mMTris-HCl, pH 8.1, 150 mMNaCl)

High salt immune complex buffer (0.1% SDS, 1% Triton X-100, 2 mM EDTA, 20 mM Tris-HCl, pH 8.1, 500 mMNaCl),

Lithium chloride immune complex buffer [0.25 M LiCl, 1% IGEPAL CA630, 1% Deoxycholic acid (sodium salt), 1 mM EDTA, 10 mMTris, pH 8.1] and

TE buffer (10 mMTris-HCl, pH 8.0, 1mM EDTA).
