## Supplementary File 4 for "MiR-146a-dependent regulation of CD24/AKT/β-catenin axis drives cancer stem cell phenotype in oral squamous cell carcinoma"

```
Seq1,Seq2,Tot Score,Tot Energy,Max Score,Max Energy,Strand,Len1,Len2,Positions
>>hsa_miR146a_5p    hg38_refGene_NM_013230  160.00  -22.84  160.00  -22.84  15653   22
1830     354
Complete

   Forward: Score: 160.000000  Q:2 to 18  R:354 to 376 Align Len (17) (88.24%) (94.12%)

   Query:    3' uugggUACCU-UAAGUCAAGAGu 5'
                     ||||| ||||||||:||
   Ref:      5' gtggaATGGAGATTCAGTTTTCa 3'

   Energy:  -22.840000 kCal/Mol


Other isoforms that are also targeted by this miRNA

>>hsa_miR146a_5p    hg38_refGene_NM_001291737   160.00  -22.84  160.00  -22.84  15652   22
1830     354

>>hsa_miR146a_5p    hg38_refGene_NM_001291738   160.00  -22.84  160.00  -22.84  15651   22
1830     354

>>hsa_miR146a_5p    hg38_refGene_NM_001291739   160.00  -22.84  160.00  -22.84  15656   22
1830     354


Homo sapiens CD24 molecule (CD24), transcript variant 5, non-coding RNA.
>>hsa_miR146a_5p    hg38_refGene_NR_117089  160.00  -22.84  160.00  -22.84  15657   22
2263     787
```
